# Systematic Evaluation of Different R-loop Mapping Methods: Achieving Consensus, Resolving Discrepancies and Uncovering Distinct Types of RNA:DNA Hybrids

**DOI:** 10.1101/2022.02.18.480986

**Authors:** Jia-Yu Chen, Do-Hwan Lim, Liang Chen, Yongli Zhou, Fangliang Zhang, Changwei Shao, Xuan Zhang, Hairi Li, Dong Wang, Dong-Er Zhang, Xiang-Dong Fu

## Abstract

R-loop, a three-stranded nucleic acid structure, has been recognized to play pivotal roles in critical physiological and pathological processes. Multiple technologies have been developed to profile R-loops genome-wide, but the existing data suffer from major discrepancies on determining genuine R-loop localization and its biological functions. Here, we experimentally and computationally evaluate eight representative R-loop mapping technologies, and reveal inherent biases and artifacts of individual technologies as key sources of discrepancies. Analyzing signals detected with different R-loop mapping strategies, we note that genuine R-loops predominately form at gene promoter regions, whereas most signals in gene body likely result from structured RNAs as part of repeat-containing transcripts. Interestingly, our analysis also uncovers two classes of R-loops: The first class consists of typical R-loops where the single-stranded DNA binding protein RPA binds both the template and non-template strands. By contrast, the second class appears independent of Pol II-mediated transcription and is characterized by RPA binding only in the template strand. These two different classes of RNA:DNA hybrids in the genome suggest distinct biochemical activities involved in their formation and regulation. In sum, our findings will guide future use of suitable technology for specific experimental purposes and the interpretation of R-loop functions.

## Introduction

R-loop is a three-stranded structure composed of an RNA:DNA hybrid and the displaced non-template DNA (Crossley et al. 2019). R-loops are thought to pose a major threat to genome stability (Gan et al. 2011; Sollier et al. 2014), underlying many human disease processes (Richard and Manley 2017). More recent studies reveal that R-loops also play regulatory roles in diverse physiological pathways, including chromosome segregation (Kabeche et al. 2018), gene expression regulation (Arab et al. 2019) and repair of DNA double-strand breaks by homologous recombination (Liu et al. 2021; Ouyang et al. 2021). These findings underscore the important impact of R-loops on many critical biological processes.

The vital biological functions of R-loops have fueled the development of two types of genome-wide R-loop mapping strategies. The first takes advantage of the specificity of the S9.6 monoclonal antibody in recognizing RNA:DNA hybrids, leading to the development of DRIP-seq (DNA:RNA Immunoprecipitation and sequencing), which sequences R-loop-containing restriction fragments enriched by S9.6 immunoprecipitation (IP) (Ginno et al. 2012). An iteration of the DRIP-seq is DRIPc-seq (DRIP followed by cDNA conversion and sequencing), designed to sequence S9.6-captured RNA (Sanz et al. 2016). Additionally, various refinements have been made (Ginno et al. 2012; El Hage et al. 2014; Chen et al. 2015; Nadel et al. 2015; Sanz et al. 2016; Wahba et al. 2016; Dumelie and Jaffrey 2017; Xu et al. 2017; Crossley et al. 2020), to name just a few, RDIP-seq (RNA:DNA IP and sequencing) by RNase I treatment and sonication followed by S9.6 IP and sequencing of RNA extension products (Nadel et al. 2015), ssDRIP-seq (single-strand DRIP-seq) by using a combination of frequent cutters to fine fragment DNA followed by sequencing S9.6-enriched DNA from the template strand (Xu et al. 2017), and bisDRIP-seq (bisulfite-based DRIP-seq), which couples S9.6-based capture with bisulfide sequencing of the displaced non-template DNA (Dumelie and Jaffrey 2017).

The second class of R-loop mapping strategies leverages a catalytically-deficient but binding-competent version of RNase H1 to capture R-loops, taking advantage of this evolutionarily-conserved enzyme in specific recognition of RNA:DNA hybrids. The first such approach developed is DRIVE-seq (DNA:RNA *in vitro* enrichment and sequencing), which utilizes purified RNase H1 to enrich for R-loop-containing restriction fragments *in vitro* for sequencing (Ginno et al. 2012). R-ChIP (R-loop chromatin IP sequencing) employs exogenously expressed RNase H1 to capture R-loops *in vivo* and then sequence the associated template DNA strand (Chen et al. 2017), and its recent derivative RR-ChIP (RNA R-ChIP) sequences RNA associated with the RNase H1 precipitant (Tan-Wong et al. 2019). To eliminate the requirement for the ectopic RNase H1 expression as with R-ChIP or RR-ChIP, MapR applies a fusion protein of RNase H1 and micrococcal nuclease (MNase) to permeabilized cells to recognize and release R-loop-containing DNA for sequencing (Yan et al. 2019). By further combing non-denaturing bisulfite chemistry, bisMapR is able to produce strand-specific R-loop profiles (Wulfridge and Sarma 2021). Finally, R-loop CUT&Tag utilizes the hybrid binding domain of RNase H1 to target Tn5 transposase to R-loop containing chromatin regions (Wang et al. 2021).

Although the development of various R-loop mapping strategies has greatly stimulated research in understanding R-loop biology *in vivo*, the results generated with different methods show some major discrepancies, so dramatically distinct that have caused grand confusion in terms of sequence feature, size, and distribution of R-loops in the genome. For example, most mapped R-loops are found to associate with GC-rich and G/C-skewed sequences (G-rich sequences in the non-template strand and C-rich sequences in the template strand) (Ginno et al. 2012; Chen et al. 2017), which is in line with early biochemical studies on RNA:DNA hybrid formation (Roberts and Crothers 1992), but at least two profiled R-loops are linked to A:T-rich sequences instead (Wahba et al. 2016; Xu et al. 2017). The average size of R-loops is ∼200-300 nucleotides in length under electron microscope (Duquette et al. 2004), but various mapped R-loops show a wide range from a few hundred nucleotides to several kilobases (Ginno et al. 2012; Sanz et al. 2016). The major discrepancy is where R-loops form in the genome. While R-loop formation at gene promoters (transcription start sites or TSSs) appears to be the general consensus, various studies also suggest the broad distribution of R-loops in gene bodies, transcription termination sites (TTSs), and even intergenic regions. Because the locations of R-loops in different genomic regions would have dramatically distinct implications in R-loop biology, it is important to systematically determine whether different R-loop mapping strategies are of specific biases in capturing R-loops in different genomic regions or how individual strategies may each introduce different types of artifacts in different degrees (Castillo-Guzman and Chedin 2021; Belotserkovskii and Hanawalt 2022).

Given the fundamental importance of these critical problems, we sought to make a systematic effort to investigate the sources of discrepancies in R-loop profiling literature. A generally accepted “gold standard” for validating mapped R-loops is their sensitivity to RNase H treatment, but such treatment before IP, as practiced in most published studies, would remove not only RNAs directly engaged in R-loops, but also those directly or indirectly associated with R-loop-engaged RNAs. In fact, our analysis suggests that different R-loop mapping strategies are each associated with a degree of shortcomings. Interestingly, our analysis also revealed a different type of RNA:DNA hybrids from typical nascent RNA-induced R-loops, which may reflect the established role of RNase H in DNA replication/repair. Our findings thus not only provide a guide for choice of suitable methods for future R-loop study, but also suggest a key criterion to differentiate R-loops involved in distinct biological pathways.

## Results

### Broad discrepancies in genome-wide R-loop profiles generated with different technologies

Eight R-loop mapping technologies were selected for comparative analysis based on their representation of different experimental strategies and resulting profiles in HEK293 cells or Hela cells, thus relatively more suitable for direct comparison (Table 1). All R-loop mapping data were processed with a uniform pipeline (see Methods). As noted previously (Halasz et al. 2017; Vanoosthuyse 2018; Crossley et al. 2019; Lin et al. 2022), different strategies have distinct performances, resulting in variable peak number (from ∼5,000 to ∼80,000) and peak size range (from ∼200 bp to ∼1.5 kb) (Fig. 1A, B). In general, S9.6-based technologies detect more peaks than RNase H-based ones (Fig. 1A). Technologies relying on DNA fragmentation by sonication usually detect sharper peaks, indicative of relatively lower mapping resolution for other technologies (Fig. 1B).

**Figure 1.**
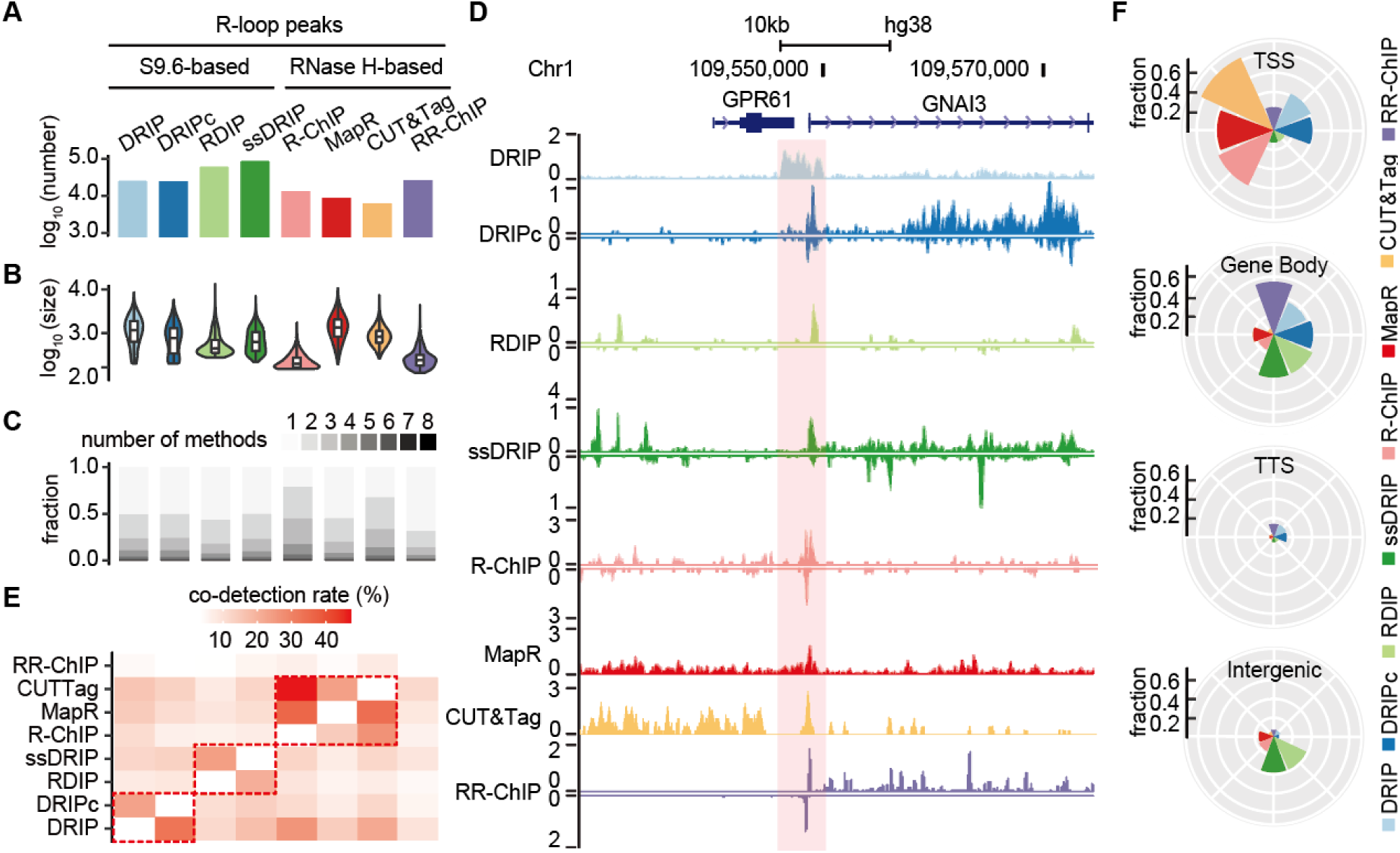
Cross comparison of R-loop mapping technologies. (A) Numbers of peaks detected with different technologies. (B) The size distributions of peaks mapped with different technologies. (C) Peaks detected with individual technologies (x-axis) are grouped based on numbers of technologies they can be detected with. For example, groups shaded in the lightest grey represent those peaks detectable with only a single technology. (D) Illustration of signals mapped by different technologies on a representative genomic locus. Highlighted in red-shaded area is a single region commonly detected by all technologies. (E) Fractions of peaks detected by each technology (x-axis) that are co-detected by other technologies (y-axis). Highlighted in red-dotted boxes are technologies that show the highest overlap. (F) The genomic annotation of detected peaks by different technologies. Here, peaks are assigned based on the following priority order: TSS > TTS > Gene Body > Intergenic Region if they are overlapped with multiple genomic regions.

**Table 1.** Overview of genome-wide R-loop mapping technologies

| Technology | Detection Method | Fragmentation Method | Molecule Sequenced | Data Resources |
| --- | --- | --- | --- | --- |
| DRIP-seq | S9.6 | restriction digestion | dsDNA | HEK293 (Manzo et al. 2018) |
| DRIPc-seq | S9.6 | restriction digestion | RNA | HEK293 (Manzo et al. 2018) |
| RDIP-seq | S9.6 | sonication | RNA | HEK293 (Nadel et al. 2015) |
| ssDRIP-seq | S9.6 | restriction digestion | ssDNA | HeLa (Yang et al. 2019) |
| R-ChIP | RNase H | sonication | ssDNA | HEK293 (Chen et al. 2017) |
| MapR | RNase H | MNase treatment | dsDNA | HEK293 (Yan et al. 2019) |
| R-loop CUT&Tag | RNase H | Tn5 tagmentation | dsDNA | HEK293 (Wang et al. 2021) |
| RR-ChIP | RNase H | sonication | RNA | HeLa (Tan-Wong et al. 2019) |

Comparative analyses were next performed to delineate the pairwise relationships for all R-loop mapping technologies after controlling for different mapping depths and cell types (see Methods). We selected a fixed set of 1-kb bins that were covered by at least one of the top 5,000 peaks scored by individual technologies for comparison. Strikingly, ∼50% of peaks were detectable by only a single technology (Fig. 1C), which is readily appreciable at a representative genomic locus (Fig. 1D). Interestingly, systematic pairwise comparison revealed three groups of technologies (i.e., DRIP-seq vs. DRIPc-seq, RDIP-seq vs. ssDRIP-seq, and R-ChIP vs. MapR vs. R-loop CUT&Tag) that show the highest overlap in peak location with each another (highlighted by dashed boxes in Fig. 1E), which likely reflects intrinsic similarity of their experimental designs. Technically, for example, both DRIP-seq and DRIPc-seq generate the same restriction fragments for S9.6 enrichment followed by sequencing S9.6-enriched DNA (DRIP-seq) or RNA (DRIPc-seq), thus explaining similar large peaks generated by this pair of technologies. RDIP-seq and ssDRIP-seq both take advantage of S9.6-enriched DNA from the template strand, either for RNA-primed extension (RDIP-seq) or adaptor ligation (ssDRIP-seq), to construct strand-specific libraries, which may produce similar peaks. Lastly, R-ChIP, MapR and R-loop CUT&Tag all leverage the specificity of RNase H1 in R-loop recognition, thus likely generating similar R-loop profiles. Unexpectedly, while highly related to R-ChIP, RR-ChIP detects a large number of peaks in gene bodies not seen with R-ChIP, and interestingly, most of those gene body peaks are often distinct from DRIPc-seq detected ones (see examples in Fig. 1D), indicating different populations of RNAs captured with RR-ChIP and DRIPc-seq (see below).

These pair-wise inter-technology relationships are also echoed in the genomic distribution of putative R-loops detected (Fig. 1F). In contrast to the remarkable enrichment for peaks at TSSs detected with R-ChIP, MapR and R-loop CUT&Tag, all S9.6-based technologies and RR-ChIP show significantly enriched signals in gene bodies, and somewhat counterintuitively, RDIP-seq and ssDRIP-seq appear to exhibit particular bias for signals in intergenic regions where transcription is less active compared to genic regions. As the genomic context is of great importance in understanding the biological functions of R-loops, these major discrepancies stress the necessity to systematically evaluate individual technologies to better understand R-loop biology and guide future research on this important aspect of functional genomics.

### Mis-assignment to genomic annotations due to low-resolution peaks

Manually examining representative broad peaks detected by DRIP-seq, we noted that the resolution limited by certain technologies tends to compromise the accurate assignment of mapped peaks in the genome. For instance (Fig. 1D), a broad DRIP-seq peak could be assigned to both the TTS of the upstream gene and the TSS of the downstream gene, while signals detected with all other technologies appear to point to the TSS region, suggesting that the TSS of the downstream gene likely corresponds to the region for R-loop formation. To address this issue more systematically, we displayed the median peak size against the percentage of peaks assignable to more than one genomic annotation with individual technologies, revealing a linear relationship, showing that the larger the median peak size is, the higher level of multi-assignment it causes (Fig. 2A). We noted two outliers with very low level of multi-assignment, i.e., RDIP-seq and ssDRIP-seq, as reads from both were largely mapped to intergenic regions (see Fig. 1F), thus having less chance for overlapping with genic regions. To quantify genomic regions that tend to cause multi-assignment, we plotted singly and multi-assigned peaks with each technology, which could be segregated into eight groups (Fig. 2B). Notably, DRIP-seq is among technologies with a very high fraction of peaks within the triple assignment group (see Supplemental Fig. S1A for a representative case).

**Figure 2.**
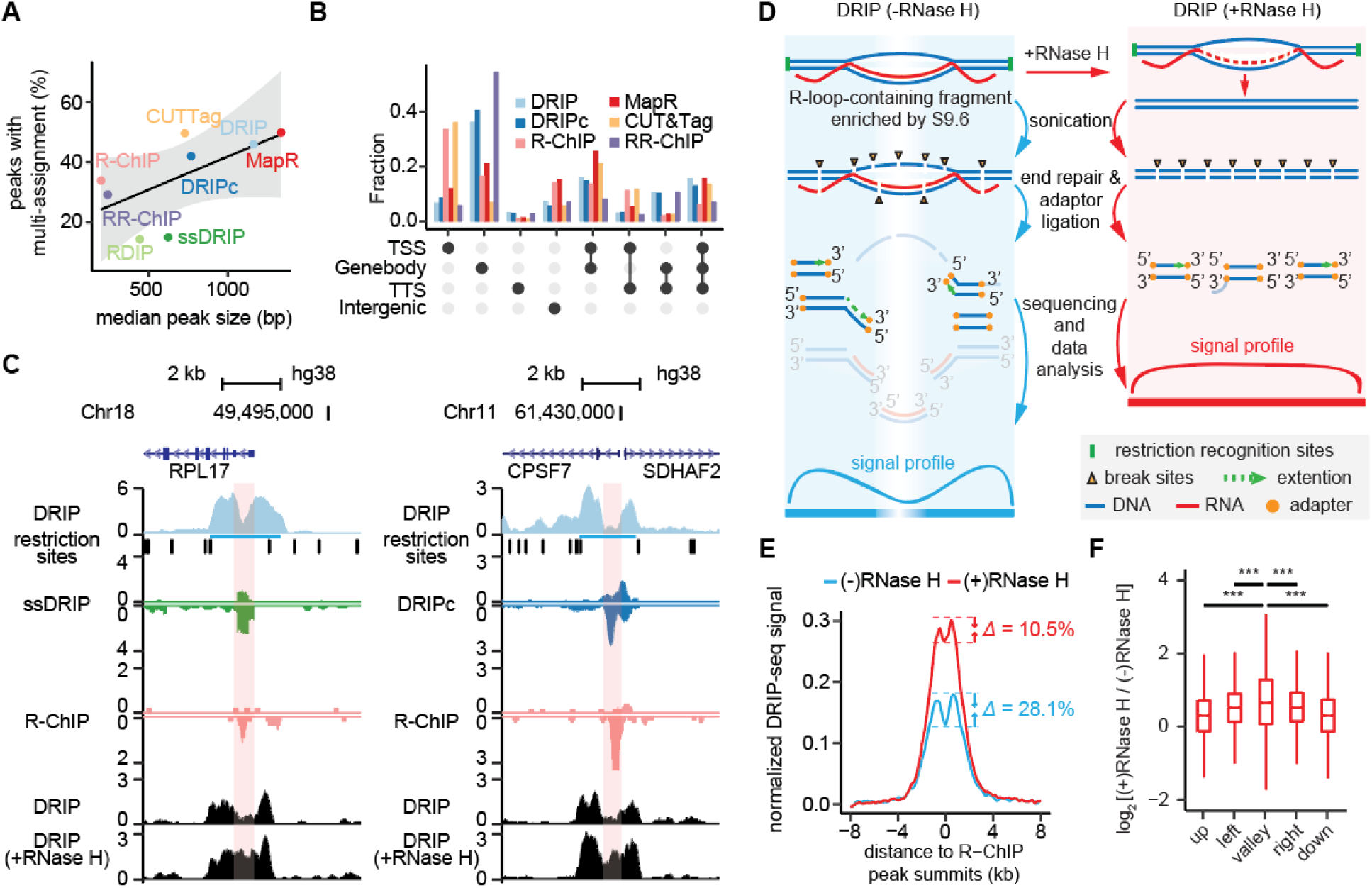
Imprecise genomic annotation of mapped peaks due to low mapping resolution. (A) Percentage of peaks assigned to multiple genomic annotations as a function of median peak size. Linear regression is presented as the regression line with standard errors of estimate (gray area). (B) The distribution of peaks assigned to different genomic annotations. Peaks are further divided into eight groups based on their assignment to one, two, or three genomic annotations. (C) Single DRIP-seq peaks at *RPL17* promoter (left) and the divergent promoter for *CPSF7* and *SDHAF2* (right) in reference to the distribution of the restriction sites and specific peaks mapped by ssDRIP-seq, DRIPc-seq and R-ChIP. The red-shaded area highlights the “valley” within a bimodal DRIP-seq peak region (light blue underlined). (D) Standard DRIP-seq procedure (left) and revised DRIP-seq by including an RNase H treatment step after S9.6 IP (right). Fragments highlighted by transparent color represent those with reduced efficiency for end-repair and/or adaptor ligation. (E) DRIP-seq signals aligned at R-ChIP peaks before and after RNase H treatment. (F) Fold-changes in DRIP-seq signal intensity in response to RNase H treatment at valleys, upstream (left) or downstream (right) of valleys, and further upstream (up) or downstream (down) regions. \*\*\**p*-value < 0.001 based on *Wilcoxon* test.

Peaks assigned to more than one genomic annotation are problematic when studying specific R-loop functions. For example, a peak associated with a TSS would implicate a role of R-loop in transcription initiation and/or pause release, while a peak with TTS would suggest a different role of such R-loop in transcription termination. Thus, identifying the source(s) for peak mis-assignment is important. We first paid special attention to DRIP-seq, the most widely-used technology with a very high level of multi-assignment (Fig. 2A, B). Theoretically, a given DRIP-seq peak should exactly correspond to the R-loop-containing restriction fragment. As anticipated, DRIP-seq peaks show enrichment of restriction sites at peak boundaries and depletion within peaks (Supplemental Fig. S1B). Unexpectedly, two separate DRIP-seq peaks are frequently observed mapped to the same restriction fragments, accounting for ∼10% of total peaks, ∼60% of which were separately assigned to different genomic annotations (see an example in Supplemental Fig. S1C). Even with the rest of restriction fragment-sized DRIP-seq peaks, many exhibit a bimodal signal distribution with a “valley” in between colocalizing with a single sharp peak detected by other technologies, such as DRIPc-seq, ssDRIP-seq and R-ChIP (Fig. 2C).

To determine how widespread this phenomenon is, we used a step detection strategy to identify regions within restriction fragment-sized DRIP-seq peaks that show a dramatic decrease in signal intensity (Supplemental Figure S2A, B; Methods). From a total of 1,907 DRIP-seq peaks that show overlap with R-ChIP peaks, for example, we could identify valleys in 93.6% (1,785) of them, about half of which (53.4%, 1,019) co-localized precisely with a single R-ChIP peak (Supplemental Fig. S2C), which is significantly higher than randomly simulated R-ChIP peaks (Supplemental Fig. S2D). Similar conclusions could be drawn when valleys in DRIP-seq peaks were compared with R-loop peaks detected by other technologies (data not shown).

### An RNase H-based strategy to aid in accurate peak assignment

To understand the basis for the bimodal DRIP-seq signals, we carefully revisited the experimental procedure for DRIP-seq and realized the potential obstruction of library construction by the presence of an R-loop (Fig. 2D, left). As S9.6-enriched R-loop-containing restriction fragments are first sonicated to generate shorter fragments for library construction (Ginno et al. 2012; Sanz and Chedin 2019), the looped single-stranded non-template DNA might be more vulnerable for breakage, and additionally, RNA hybridized on the template DNA might also interfere with end repair and/or adaptor ligation. As a result, the DNA within the R-loop might have reduced representation relative to the flanking regions in the resulting library.

To experimentally test this possibility, we implemented an RNase H treatment step after (rather than before, as with the common R-loop mapping validation approach) S9.6 IP and before sonication, generating overall similar DRIP-seq libraries under both experimental conditions (Supplemental Fig. S2E). This treatment would remove the RNA moiety within RNA:DNA hybrids to allow re-annealing of separated DNA strands (Fig. 2D, right), thus helping more even representation of DNA fragments in the R-loop forming region during library construction, thereby “rescuing” signals in the valley. Indeed, we found that DRIP-seq signals at valley regions significantly increased in response to RNase H treatment, as illustrated on the representative cases (Fig. 2C). Genome-wide, although smaller valleys were still detectable likely due to insufficient RNase H treatment, those valleys became shallower from 28.1% to 10.5% of peak height (Fig. 2E). It is also evident that RNase H treatment significantly elevated the signals toward the valley relative to the surrounding up- and downstream regions (Fig. 2F), which may result from multiple smaller R-loops that collectively contributed to the large peak. These results suggest a revised DRIP-seq strategy based on recovered signals after RNase H treatment, which is particularly important for accurate peak assignment to TSS or TTS of adjacent genes given their prevalence in the human genome.

Broad peaks are more prominent in gene bodies, which has been an open question with respect to frequent R-loop formation during transcription elongation (see a theoretical consideration of this issue in Discussion) (Castillo-Guzman and Chedin 2021; Belotserkovskii and Hanawalt 2022). A study in fission yeast suggests that most genic DRIPc-seq signals result from dsRNA, rather than genuine R-loops, because S9.6 was found to have significant affinity for dsRNA, and the majority of gene body signals could be eliminated by treatment with dsRNA-specific RNase III (Hartono et al. 2018). Interestingly, besides S9.6-based methods, RR-ChIP, which sequences RNA bound to exogenously expressed RNase H1 (Tan-Wong et al. 2019), also detected an alarmingly high level of signals in gene bodies, while R-ChIP, a highly related protocol that sequences RNase H1 bound DNA, detected little such signals (see Fig. 1F). This is particularly evident when we intersected peaks identified previously by DRIP-seq versus DRIPc-seq (both S9.6-based) (Manzo et al. 2018) and those captured with R-ChIP versus RR-ChIP (both RNase H-based) (Fig. 3A), finding that peaks uniquely identified by either DRIP-seq or DRIPc-seq (Fig. 3A, left) and peaks exclusively detected with RR-ChIP (Fig. 3A, right) are predominately mapped to gene bodies.

**Figure 3.**
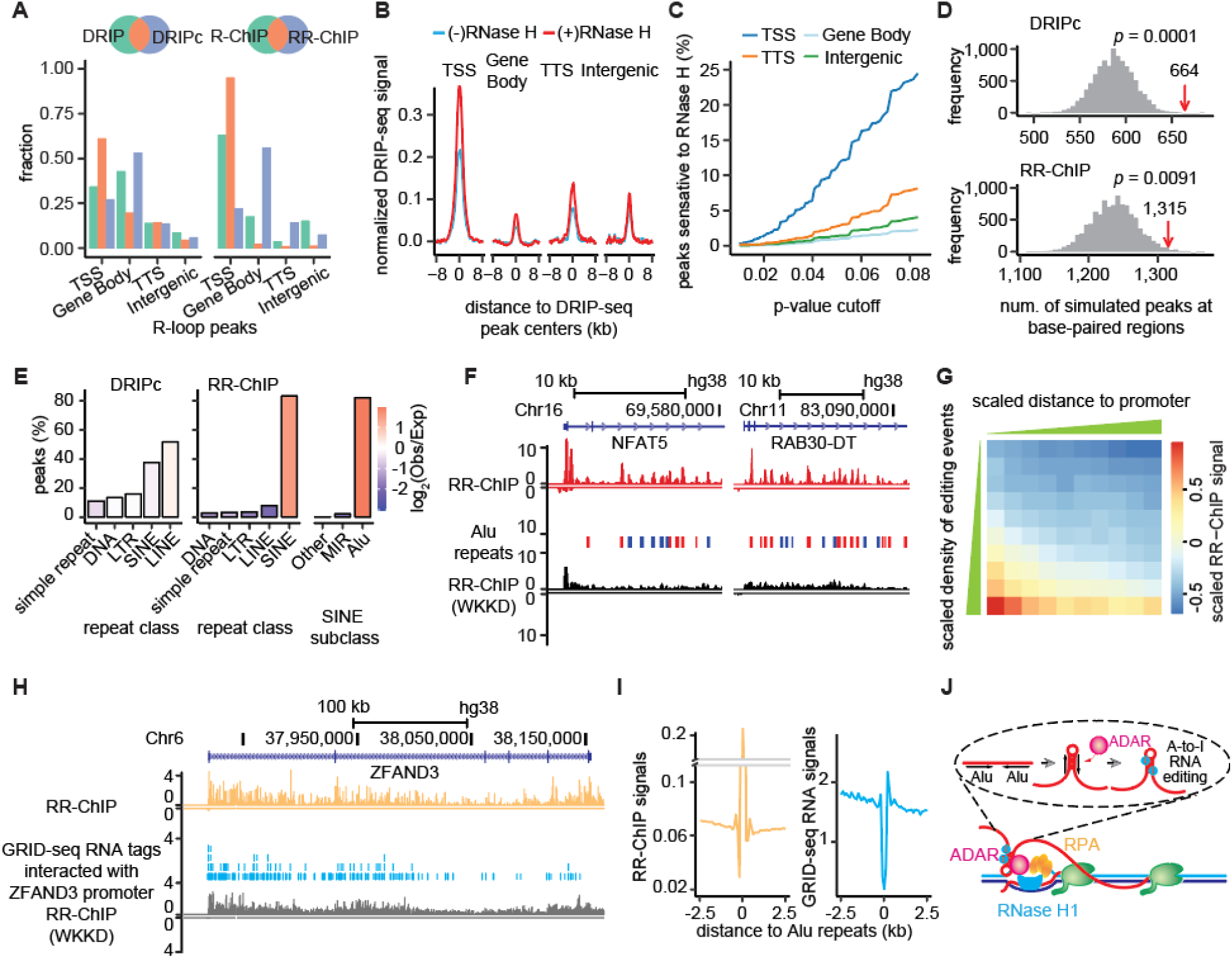
Chromatin-associated structured RNAs contribute to false-positive genic R-loop signals. (A) Genomic distribution of peaks detected by DRIP-seq (green), DRIPc-seq (blue) and both (red), and those detected by R-ChIP (green), RR-ChIP (blue) and both (red). (B) Signal profiles before (blue) and after (red) RNase H treatment in different genomic regions. (C) Percentages of DRIP-seq peaks that are sensitive to RNase H treatment across different p-value cutoffs. (D) Distribution of the expected frequency of overlap between DRIPc-seq (top) or RR-ChIP (bottom) peaks and RNA duplex regions detected by PARIS technology after 10,000-time simulations. Red arrow marks the observed overlap. *P*-values were determined as the likelihood of events with no less than the observed value. (E) Association of DRIPc (left) or RR-ChIP (right) peaks with different classes of repeats annotated by RepeatMasker. (F) RR-ChIP signals at the *NFAT5* and *RAB30-DT* gene loci in reference to the background detected with RR-ChIP (WKKD) and the distribution of Alu repeats. Alu elements on Watson and Crick strands are coded in red and blue, respectively. (G) Segregation of Alu elements into 10×10 groups based on the propensity of forming structured RNAs measured by the density of RNA editing events (columns) and their distance to promoter regions (rows). For each group, median RR-ChIP signal is shown and color-coded according to the key on the right. (H) A representative case with RR-ChIP signals and GRID-seq RNA tags interacting with the *ZFAND3* promoter. (I) Aligned on the center of Alu elements in gene bodies are RR-ChIP captured RNAs (left) in comparison with RNAs linked to gene promoters detected by GRID-seq (right). (J) A putative model depicting the origin of RR-ChIP-mapped signals on Alu elements.

We reasoned that if gene body signals were from genic R-loops, the signals would become enhanced in response to RNase H treatment after S9.6 IP but before DNA fragmentation. We thus utilized our newly generated DRIP-seq data before and after RNase H treatment to evaluate the RNase H sensitivity of peaks assigned to different genomic annotations. Meta-gene analysis of all DRIP-seq peaks showed that TSSs contained most gained signals relative to other genomic annotations (Fig. 3B). We next assessed whether individual DRIP-seq peaks showed significantly gained signals upon RNase H treatment to determine the fraction of RNase H sensitive peaks assigned to different genomic annotations (see Methods). We found that across all *p*-value cutoffs, the number of peaks that showed gained signals upon RNase H treatment is the highest at TSSs, modest at TTSs, and lowest in gene bodies and intergenic regions (Fig. 3C). Interestingly, the majority (∼70% at the cutoff of *p*-value < 0.05) of those RNase H sensitive peaks belonged to those commonly detected by DRIP-seq and DRIPc-seq, implying that most peaks uniquely detected with DRIP-seq or DRIPc-seq unlikely result from R-loop formation. These data strongly suggest that R-loops predominantly form at TSSs and with much reduced frequency in other genic regions.

### Repeat-containing structured RNAs in gene bodies co-purified with S9.6 and RNase H1

We next sought to understand the nature of RNAs captured with DRIPc-seq and RR-ChIP. Given the identification of RNAs sensitive to the treatment of RNase III, a dsRNA-specific nuclease, in fission yeast (Hartono et al. 2018), we intersected DRIPc-seq peaks in gene bodies with RNA duplex regions in HEK293 cells mapped with PARIS, a technology specifically designed to detect RNA secondary structures (Lu et al. 2016), and found that peaks detected with the RNA-based DRIPc-seq technology showed a significant overlap with mapped dsRNA regions compared to randomly simulated ones (Fig. 3D; Methods). Interestingly, peaks identified by another RNA-based RR-ChIP technology also exhibited a similar degree of overlap with structured RNAs mapped in Hela cells by PARIS (Fig. 3D). This prompted us to determine the nature of RNAs enriched with both DRIPc-seq and RR-ChIP. We found that DRIPc-seq captured signals predominately corresponded to various repeat-derived RNAs, particularly LINEs and SINEs, whereas RR-ChIP enriched signals were mainly from the Alu subfamily of SINE repeats (Fig. 3E). This is evidenced on two representative genes where individual RR-ChIP peaks are largely coincident with Alu repeats in the gene body, which were essentially absent in RR-ChIP control experiment with the hybrid binding deficient quadruple mutant (WKKD) RNase H1 (Fig. 3F).

We next wished to understand why R-ChIP (DNA-based) mainly detects R-loops at TSSs while RR-ChIP (RNA-based) additionally captures Alu RNAs in gene bodies. As implied by a recent study (Bai et al. 2021), Alu RNAs in gene bodies might form *trans*-acting R-loops at Alu-containing TSS regions given the overall sequence similarity of Alu elements. If this is true, given spatial proximity, an Alu RNA most likely would target its own TSS transcribing an Alu element of the same directionality. However, we noted that most (∼90%) of TSS R-loops do not contain Alu elements (Fig. 3F, left), and Alu RNAs in either same or opposite direction are enriched by RR-ChIP in a comparable level (Fig. 3F), likely disfavoring the formation of R-loops in *trans* by Alu RNAs in gene bodies. However, we cannot fully exclude the possibility that these Alu RNAs may form *trans*-acting R-loop at TSS regions of other genes.

Another possibility is that RNase H1 might directly bind Alu-containing transcripts, as RNase H1 has been shown to have a degree of affinity of dsRNA (Nowotny et al. 2008). Alternatively or additionally, over-expressed RNase H1 in RR-ChIP experiment may interact with Alu RNA binding proteins, thereby bringing Alu RNAs to the proximity of gene promoters with R-loop formation. In agreement with this possibility, RNase H1 has been reported to physically interact with ADAR (Rouillard et al. 2016), which can bind Alu elements to catalyze A-to-I editing (Nishikura 2010), and ADAR is also capable of binding RNA:DNA hybrids (Zheng et al. 2017; Jimeno et al. 2021), and Z-DNA structures, which are prevalently distributed at gene promoter regions (Shin et al. 2016). To obtain initial evidence for this intriguing possibility, we divided Alu elements into 10 bins based on their distances to gene promoters and 10 bins based on the density of recorded A-to-I editing events (Picardi et al. 2017). We observed that RR-ChIP signals in gene bodies were positively correlated with the density of A-to-I editing events and negatively with distance to gene promoters (Fig. 3G; Supplemental Fig. S3).

Through either direct or indirect mechanisms above, a critical prediction is that Alu-containing regions in nascent transcripts might be looped to the proximity of TSSs, thus showing higher frequencies in association with gene promoters compared to flanking regions in the same transcripts. To test this idea, we leveraged our recent data generated with the GRID-seq technology designed to detect global RNA-DNA interactions (Li et al. 2017), which allowed us to compare levels of RNAs from different transcript regions that are linked to gene promoters through proximity ligation. Indeed, this appears to be the case on individual genes (Fig. 3H). Genome-wide, we aligned the center of Alu elements with RR-ChIP reads in gene bodies (Fig. 3I, left) in comparison with GRID-seq RNA reads that were linked to promoter DNA (Fig. 3I, right), both showing a dip (which likely corresponds to the mapping gap within Alu repeats) followed by a peak. These findings provide supporting evidence that RR-ChIP captured Alu RNAs in gene bodies result from their association with gene promoters (Fig. 3J).

Collectively, our analysis suggests that most gene body signals captured with either DRIPc-seq or RR-ChIP likely result from structured RNAs, rather than genuine R-loops.

### A signification fraction of intergenic signals lack evidence for nascent RNA production

Similar to gene body signals, peaks in intergenic regions also showed limited sensitivity to RNase H treatment (see Fig. 3B), which prompted us to investigate the nature of those intergenic signals, particularly those prevalently identified by RDIP-seq and ssDRIP-seq. It is conceivable that specific intergenic regions may harbor transcription enhancers, activation of which might thus induce the formation of R-loops. We therefore asked whether peaks mapped with different technologies were associated with nascent RNA production, which has been extensively profiled in both HEK293 and HeLa cells by global run-on sequencing (GRO-seq) (Chen et al. 2017; Fei et al. 2018). We found that most peaks detected with DRIP-seq, DRIPc-seq, R-ChIP, R-loop CUT&Tag, MapR, and RR-ChIP were associated with nascent RNA production, and given that DRIPc-seq, R-ChIP, and RR-ChIP also offer strand-specific information, we further noted that the sequenced strand was largely in line with the orientation of the RNA moiety in R-loops (Fig. 4A, first panel). However, ∼30% of R-ChIP detected peaks, and more dramatically, ∼70% of peaks captured with either RDIP-seq or ssDRIP-seq had little evidence for nascent RNA transcription (Fig. 4A, first panel). When separately analyzed on different genomic annotations, all technologies detected the highest degree of association with nascent RNA production at TSSs (Fig. 4A, second panel). In contrast, RDIP-seq, ssDRIP-seq, and R-ChIP each exhibited much reduced association with nascent RNA production in gene bodies as well as at TTSs, ∼30 to 50% of which were even in the opposite orientation to RNA-engaged R-loops, and most significantly, the fraction of peaks linked to nascent RNA production went down to ∼5 to 20% in intergenic regions (Fig. 4A, third to fifth panels).

**Figure 4.**
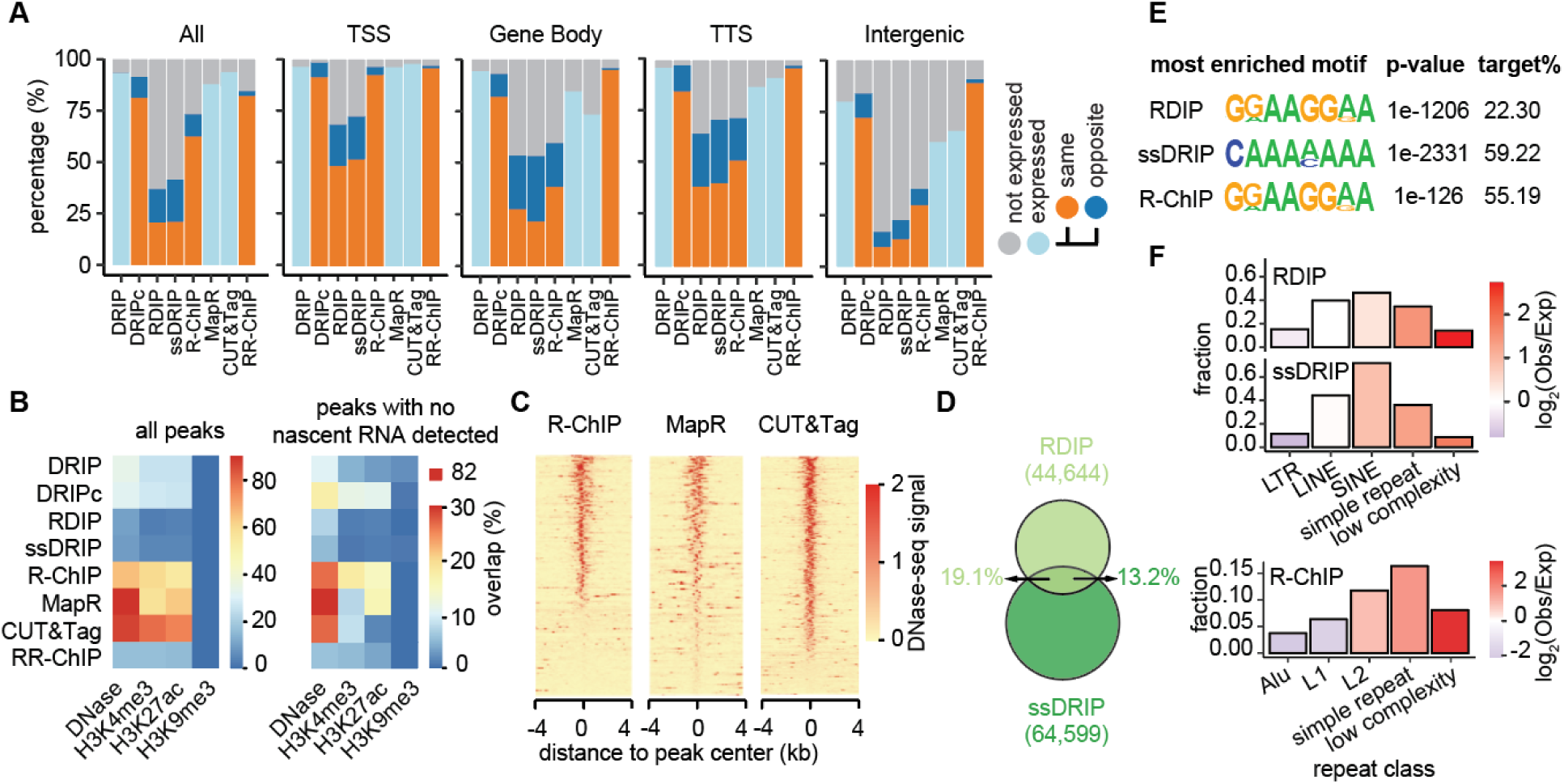
R-loop peaks lack of nascent RNA transcription are associated with open chromatin regions or simple repeats. (A) Percentages of peaks detected with different technologies without (grey) or with (other colors) evidence for nascent RNA production. For libraries without strand information, nascent RNAs from both strands are considered (light blue). For strand-specific libraries, nascent RNAs transcribed from the template (red) or non-template (dark blue) DNA are shown separately. (B) Percentages of all peaks (left) or peaks without nascent RNAs (right) associated with DNase I hypersensitivity hotspots or specific histone modification events. (C) Association of R-ChIP, R-loop CUT&Tag and MapR-mapped peaks with DNase I-seq signals. (D) Intersection of peaks detected with RDIP-seq and ssDRIP-seq with Venn diagram. (E) Most enriched 8-mer sequence motifs for RDIP-seq and ssDRIP-seq peaks, and R-ChIP peaks with no evidence of nascent RNA transcription or active histone modifications. (F) Percentages of peaks located at different repeat classes. Color key indicates the frequency of repeat-associated peaks (observed) over the expected frequency based on randomly-simulated peaks (expected).

To rule out the possibility that certain enhancer-produced RNAs (eRNAs) in intergenic regions might be below the detection limit by GRO-seq, we also examined the association between these putative R-loop forming regions with specific chromatin markers (Fig. 4B). R-ChIP, R-loop CUT&Tag and MapR mapped peaks were strongly associated with open chromatin region (DNase-seq) and active histone marks for promoter (H3K4me3) and enhancer (H3K27ac) while depleted at heterochromatin regions (H3K9me3). This was true on all detected peaks (Fig. 4B, left) or on peaks without detectable GRO-seq signals (Fig. 4B, right). In contrast, RDIP-seq and ssDRIP-seq mapped peaks likely lack such association (Fig. 4B), suggesting that many of their enriched peaks do not correspond to nascent RNA-induced R-loop formation.

Of note, R-loop CUT&Tag-mapped peaks are likely more associated with broader and stronger DNase-seq peaks (Fig. 4C), suggesting the relatively higher sensitivity of R-loop CUT&Tag method to R-loops at more opened chromatin regions. Literally, all MapR-mapped peaks colocalized with DNase I hypersensitive hotspots, including those without evidence for transcription (Fig. 4C). This indicates that the MNase-RNase H1 fusion protein may attack open chromatin regions, even in the absence of R-loops, despite that MapR signals are detected against control libraries generated with MNase alone (Yan et al. 2019). This suggests a necessity to develop more robust experimental protocol (Jauregui-Lozano et al. 2022) and statistical model to infer MapR-mapped R-loops in reference to the background with MNase treatment.

### Enriched repeat sequences in non-transcribing regions

We next wished to understand the sequence features of RDIP-seq and ssDRIP-seq enriched peaks lacking evidence of nascent RNA transcription. We first noted that there was limited overlap between peaks in non-transcribing regions detected with these two technologies (Fig. 4D), despite the fact that these two technologies are overall more related to each other than to others (see Fig. 1E). Through analyzing the motifs underlying these peaks, we found that the top motif enriched by RDIP-seq is a simple repeat sequence containing GA dinucleotides whereas that selected with ssDRIP-seq is a low-complexity A-rich sequence, which are present in 22.3% and 59.2% of total peaks, respectively (Fig. 4E). This reminded us of extensive repeats of different classes associated with the peaks enriched by DRIPc-seq and RR-ChIP (see Fig. 3E). Indeed, both RDIP-seq and ssDRIP-seq detected peaks were prevalently associated with LINE, SINE, simple repeats, and low complexity sequences, the latter two of which were more enriched compared to their relative representations in the genome (Fig. 4F).

Although both RDIP-seq and ssDRIP-seq use S9.6 for enrichment, we suspected that the RDIP-seq specific bias might result from the selection of ssDNA with simple repeat sequences after sonication and annealing of reverse complement ssDNA from the opposite strand to initiate primer extension during library construction. On the other hand, during the ssDRIP-seq procedure, abundant low-complexity sequences might have higher tendency to expose ssDNA regions that are selectively built into the library. Since both RDIP-seq and ssDRIP-seq are designed to target RNA:DNA hybrids, this raises an intriguing possibility that certain RNA components may be associated with at least a fraction of those simple repeats or low complexity sequences, even though those RNAs might not be nascent transcripts generated by the engaged RNA polymerase in the GRO-seq experiment. Notably, R-ChIP-mapped peaks lack of nascent RNA transcription or active histone modifications are also significantly associated with simple repeats or low complexity sequences (Fig. 4E, F). The most enriched motif of these R-ChIP-mapped peaks is exactly the same as that of RDIP-seq (Fig.4E). All above observations began to suggest the existence of a different type of RNA:DNA hybrids enriched by various R-loop mapping strategies (see below).

### Independent evidence for the formation of R-loops

Given the diverse peak profiles revealed by different R-looping mapping technologies, we next sought for independent evidence to evaluate the performance of different technologies. Replication protein A (RPA) is a primary candidate for this purpose (Garcia-Muse and Aguilera 2019). RPA is a heterotrimeric complex consisting of RPA1, RPA2, and RPA3 subunits, which is a major ssDNA binding protein that has been long implicated in DNA replication/repair pathways (Marechal and Zou 2015). Interestingly, RPA has been recently reported as part of R-loops where it binds the displaced non-template DNA via protein-DNA interactions as well as RNase H through direct protein-protein interactions, which is thought to help RNase H recruitment *in vivo* and stimulate R-loop resolution (Nguyen et al. 2017). This observation suggests that the binding profile of RPA may provide independent evidence for R-loop formation in the genome.

To test this idea, we first utilized a public RPA ChIP-seq dataset (Zhang et al. 2017) to compare with peaks detected with different R-loop mapping strategies and found varying degree of overlaps, highest with R-ChIP (Fig. 5A, left). Since the existing RPA ChIP-seq dataset only identified 987 peaks, and there was no strand information for its binding, we generated our own RPA ChIP-seq libraries on HEK293 cells by using a strand-specific strategy during library construction (Supplemental Fig. S4A; Methods). With the highly reproducible libraries (Supplemental Fig. S4B), we identified 2,559 significant peaks, and although the number of identifiable peaks was still lower than those detected with various R-loop mapping strategies, we observed much higher overlaps as compared to the previous dataset, again highest with R-ChIP (Fig. 5A, right). Importantly, the coincident peaks occurred not only on TTSs, as expected, but also on discrete intergenic regions, as seen on a representative genomic locus (Fig. 5B).

**Figure 5.**
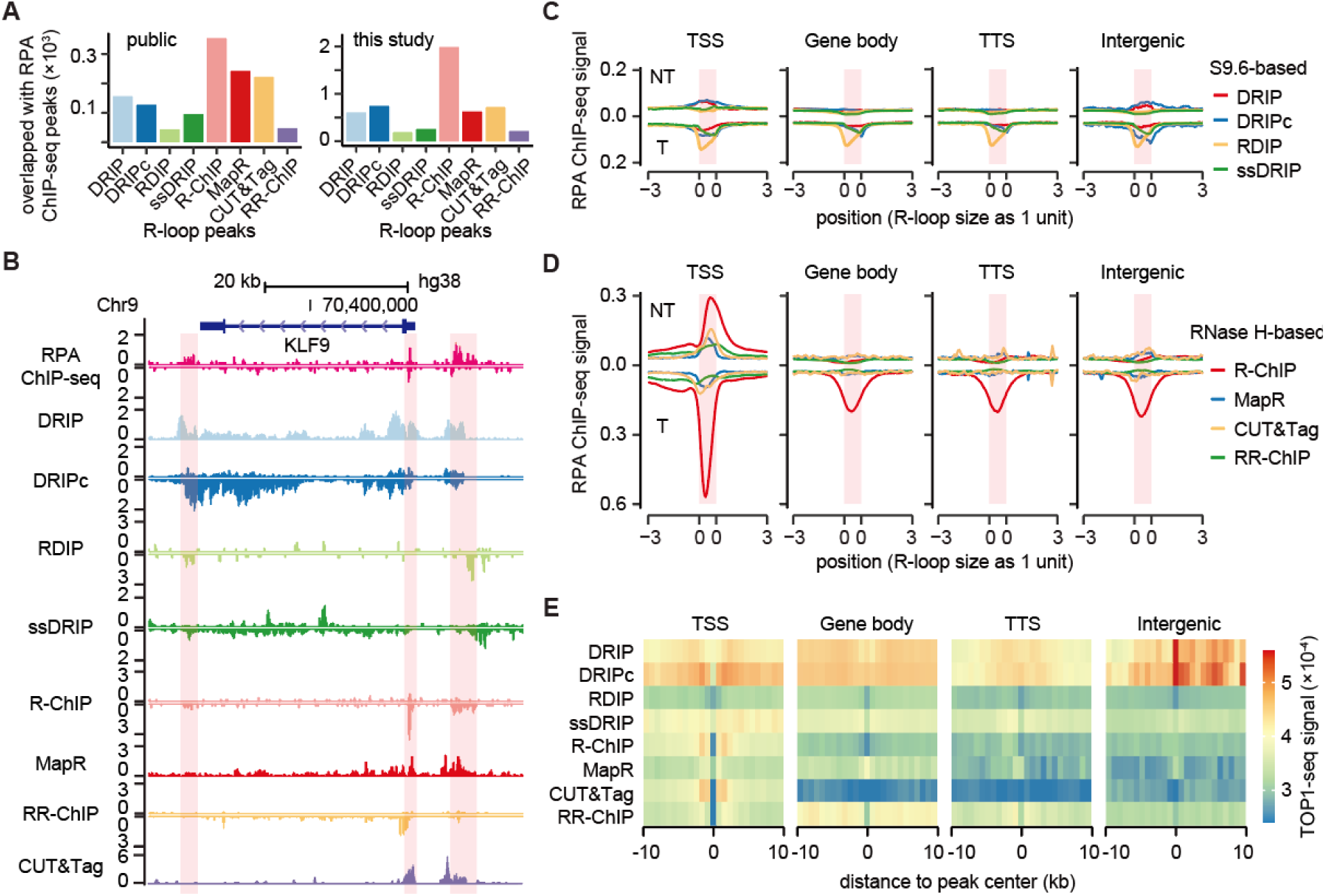
Assessing R-loops peaks with independent approaches. (A) Number of R-loop peaks that colocalize with RPA binding sites identified in a previous study (left) (Zhang et al. 2017) and current study (right). (B) Comparison of RPA binding with signals detected with different R-loop mapping technologies on a specific genomic locus. Red-shaded areas highlight three RPA binding regions. (C and D) Alignment of strand-specific RPA binding signals on peaks identified with S9.6-based technologies (C) and with RNase H-based technologies (D) on different genomic annotations. NT: non-template strand; T: template strand. (E) Meta-gene analysis to compare TOP1 activity measured by TOP1-seq with peaks detected by individual R-loop mapping technologies at different annotated genomic regions.

We next performed meta-gene analysis separately on different genomic annotations. In this analysis, we took advantage of the strand-specific information on RPA binding from the newly mapped dataset, and by aligning peaks mapped by different R-loop mapping methods, we found that all S9.6-based technologies showed the modest association of RPA binding signals (Fig.5C) compared to RNase H-based technologies (Fig. 5D) at all genomic annotations. Among the four RNase H-based technologies, R-ChIP showed the most robust correlation with the mapped RPA peaks. Focusing on R-ChIP mapped peaks in different genomic annotations, we noted that TSSs showed RPA binding on both the non-template (NT) and template (T) strands, with the peak on the non-template DNA slightly shifted to the 3’ end (Fig. 5D, first panel), which is likely due to the formation of structured DNA in the non-template strand to aid in DNA opening for RNA invasion into bubbled DNA to form R-loops. Strong RPA binding on the template strand is somewhat unexpected as RPA is generally thought to bind the displaced non-template ssDNA in R-loops. It is not due to antisense R-loop formation by prevalent divergent transcription at TSS regions because the pattern persists even when we excluded bidirectional R-loop regions (Supplemental Fig. S4C). We suspected that the strong association with the template strand likely resulted from the association of RPA with RNase H, as reported earlier (Nguyen et al. 2017). It is also possible that RPA might displace a portion of nascent RNA annealed to the template DNA, thus occupying exposed ssDNA regions during R-loop resolution. Unexpectedly, RPA appears to predominantly bind the template strand in R-ChIP identified peaks in gene bodies, TTSs, and intergenic regions (Fig. 5D, the second to fourth panels). This provides supporting evidence for the presence of two distinct types of RNA:DNA hybrids, distinguishable by the RPA binding patterns, which is also evident, although to much less extent, with various S9.6-based technologies (see Fig. 5C).

As part of our efforts in searching for independent evidence to verify the formation of genuine R-loops in the genome, we also came across an activity map for topoisomerase I (Top I), which was generated by sequencing DNA ends upon trapping the TOP1-DNA cleavage complex with Camptothecin (Baranello et al. 2016). Top I is well known to release negative supercoiling to counteract non-B DNA structure formation, and conversely, stable non-B DNA structure may suppress Top I activity. When encountering genomic regions prone to R-loop formation, such transient suppression may create a time-window and spatial proximity for nascent RNA to invade into dsDNA. We thus asked whether local suppression of TOP1 activity might be associated with R-loop formation across the genome. As the scarcity of mapped TOP1 activities only permitted meta-gene analysis, we aligned peak centers identified by different R-loop mapping technologies in reference to mapped TOP1 activities (Fig. 5E; Methods). On different genomic annotations, we observed that TOP1 activity was indeed suppressed at TSS-associated peaks identified by all published R-looping technologies, although modest for DRIP-seq and DRIPc-seq. We also observed modest repression of TOP I activity in gene bodies, TTSs, and intergenic regions with the exception of the peaks detected by MapR, and for unknown reason, even enhanced TOP I activity in intergenic peaks identified by DRIP-seq and DRIPc-seq. Although insufficiently definitive, the strongest negative association with TOP1 activity supports R-loop formation predominately at TSSs compared to other genomic regions.

### RPA binding profile and local transcription differentiate two classes of RNA:DNA hybrids

The observation of RPA binding on both the template and non-template strands of DNA at TSSs, but only the template strand in gene bodies, TTSs, and intergenic regions (see Fig. 5D) provides a critical criterion to differentiate between two distinct types of RNA:DNA hybrids in the genome. In fact, RPA is best known for its roles in DNA replication/repair pathways (Marechal and Zou 2015), which involve not only RNA, but also RNase H (Ohle et al. 2016; Hawley et al. 2017; D’Alessandro et al. 2018; Bader et al. 2020). This raises the possibility for the presence of RNA:DNA hybrids associated with DNA replication/repair events, which may actually correspond to signals in specific genomic regions that do not have evidence for typical RNA polymerase-catalyzed nascent RNA production.

Given the strongest association of R-ChIP mapped peaks with RPA binding in all genomic annotations (see Fig. 5D), we focused on comparison of R-ChIP generated peaks with RPA binding and transcription of nascent RNA detected by GRO-seq. Strikingly, we found that R-ChIP detected peaks could indeed be segregated into two separate classes, one associated with nascent RNA production and the other without detectable GRO-seq signals (Fig. 6A, left two panels). Interestingly, the first class associated with nascent RNA production showed symmetric RPA binding to the template DNA strand centered at R-ChIP peaks, but asymmetric RPA binding to the non-template DNA strand on only the downstream side; in contrast, the second class without nascent RNA synthesis exhibited only symmetric RPA binding to the template DNA strand centered at the peaks (Fig. 6A, right two panels). A similar observation was made when R-ChIP generated peaks were classified based on actively transcribed Pol II RNAs detected by NET-seq, a nascent RNA mapping technology with higher resolution (Fig. 6B) (Mayer et al. 2015). Based on these observations, we suggest that the first class corresponds to genuine R-loops (Fig. 6C, top) whereas the second class likely represents RNA:DNA hybrid formation at DNA replication forks, which may enlist primase-generated RNA (Baranovskiy et al. 2016). Alternatively, this second class of RNA:DNA hybrids may also result from transcription on single-stranded DNA at DNA damage sites where the non-template DNA is resected during the repair process, and RNA polymerase is recruited to the template DNA to generate DNA damage-induced long non-coding RNA known as dilncRNA (Hawley et al. 2017; Bader et al. 2020; Liu et al. 2021) (Fig. 6B, bottom). The greatly reduced stability of RNA polymerase on ssDNA compared to typical dsDNA, as observed with a bacterial RNA polymerase (Zenkin et al. 2006), provides a potential explanation to the lack of RPA binding to the non-template DNA and detectable nascent RNA signals by GRO-seq or NET-seq, which requires the stable association of paused RNA Polymerase on DNA to resume transcription (see Discussion). In support of the potential association between the Class 2 hybrids and DNA damage repair, we found that Class 2 hybrids are more located within late replicating regions, which are associated with a higher rate of mutation as a direct result of DNA damage (Stamatoyannopoulos et al. 2009). Regional variation of replication timing has been mapped via Repli-seq (Hansen et al. 2010). As expected, we noted that Class 2 hybrids show stronger association than Class 1 with late S phase replication domains that are conservative across different cell lines (Fig. 6D). Despite of all the above evidence, future studies are clearly needed to comprehensively unravel the nature and functions of these two distinct classes of hybrids.

**Figure 6.**
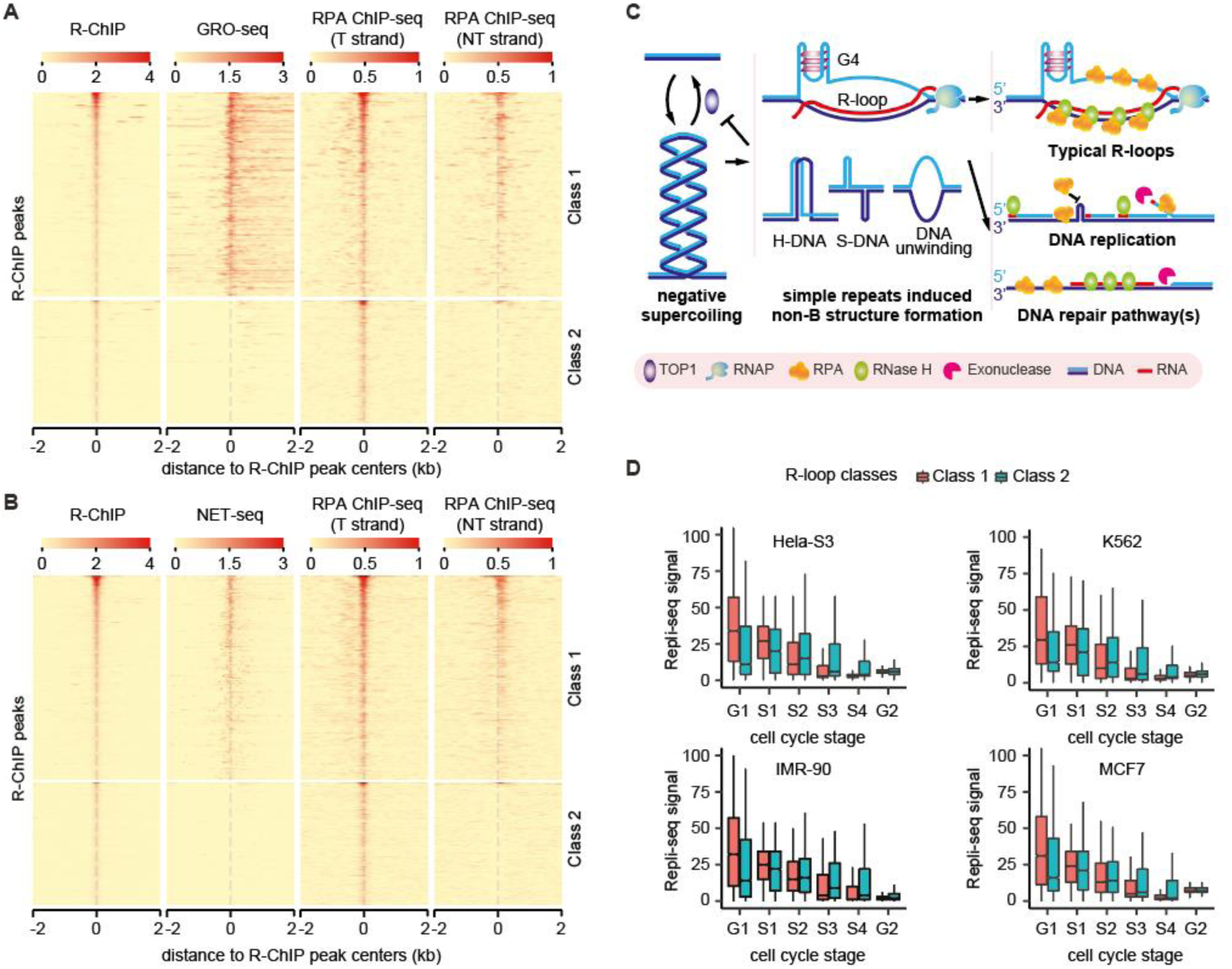
Two types of RNA:DNA hybrids and their functional connections with RPA and TOP1. (A and B) Signal profiles of R-ChIP, GRO-seq, and RPA ChIP-seq are shown for R-ChIP peaks with (Class 1) or without (Class 2) nascent RNA production detected by GRO-seq (A) or NET-seq (B). (C) Negatively supercoiled DNA is prone for the formation of various non-B structures if not relaxed by topoisomerase TOP1, and non-B structure may in turn suppress TOP1. At classical R-loop regions, RPA binds directly to the single-stranded non-template DNA, as well as the template strand through protein-protein interactions with RNase H1 to stimulate R-loop resolution. RPA also plays important roles in DNA replication, including counteracting slipped structure formation of single-stranded repeat sequences or protecting flipped ssDNA. Persistent non-B DNA may cause genome instability. In many DNA repair pathways, resection of the other strand by 5’ exonuclease generates ssDNA, which can be recognized by RPA. In both DNA replication and repair pathways, RNase H1 as well as RNase H2 have been suggested to process RNA:DNA hybrids formed by RNA primer or damage-induced RNA. (D) The distribution of Repli-seq signal at Class 1 and Class 2 RNA:DNA hybrids in four cell types.

## Discussion

### Choice of methods for R-loop mapping in the genome

Our current study was initially motivated to address the alarmingly high discrepancy among different R-loop profiling technologies in order to provide a guide for the choice of suitable methods in future studies of R-loop biology. S9.6-based methods all leverage the high binding affinity of the antibody for RNA:DNA hybrids. However, there are two main problems associated with these approaches. The first is the limited resolution. Isolated RNA-containing genomic DNA has to be fragmented before S9.6 IP. If sonication without cell fixation is used for this purpose, certain fragile R-loops may be destroyed during the process. It appears that restriction digestion with multiple frequent cutters is the most optimal strategy, as with ssDRIP-seq, which has the additional advantage in generating strand-specific library (Xu et al. 2017). The second problem is that S9.6 also has a considerable affinity for other forms of RNA and perhaps DNA. Various refinements have been proposed or developed to eliminate contaminated RNAs by using RNase T1 or RNase I for ssRNA and RNase V1 or RNase III for dsRNA (Nadel et al. 2015; Hartono et al. 2018; Sanz and Chedin 2019). However, these approaches are not widely put into practice, and cannot eliminate non-B DNA structures formed by repetitive sequences that might be also captured by the antibody, as suggested by our current findings. A key criterion for validating R-loops is to identify RNase H-sensitive peaks by generating libraries before and after RNase H treatment. However, as RNA directly or indirectly attached to R-loop engaged RNA would be also removed, caution must be taken to accurately interpret the results.

RNase H-based methods are also associated with various limitations. The use of purified RNase H to enrich for RNA:DNA hybrids as with DRIVE-seq appears to suffer from insufficient binding affinity (Ginno et al. 2012), probably because of missing enhancing factors in the *in vitro* binding reaction. This also applies to MapR (Yan et al. 2019), and possibly bisMapR (Wulfridge and Sarma 2021), as indicated by the much fewer peaks captured. In addition, MapR is also interfered with non-specific actions of fused MNase on DNase I hypersensitivity sites despite the use of MNase-treated samples as control, which could be efficiently mitigated in bisMapR by further coupling non-denaturing bisulfite chemistry. R-loop CUT&Tag (Wang et al. 2021) is very appealing and has our strong recommendation, but might also have a higher tendency to map R-loops within strong DNase I hypersensitive hotspots. R-ChIP (Chen et al. 2017; Chen et al. 2019) seems to produce the most robust peaks to date. However, the requirement for expressing exogenous RNase H1 is clearly a major hurdle in many applications. Additionally, exogenous catalytically dead RNase H1 may artificially alter the dynamics of R-loops by competing with endogenous enzymes, which may explain the significant enrichment of R-ChIP signals at TSS regions that are stabilized during the transcription cycle (Tan-Wong et al. 2019). RR-ChIP is the latest extension of R-ChIP (Tan-Wong et al. 2019), but it is associated with a different problem due to highly enriched Alu repeat-containing transcripts. Therefore, none of the technologies developed to date is perfect in every aspect. Future users have to carefully consider the pros and cons of individual technologies, filter out technology-specific false positive signals, and study the RNA:DNA hybrid class of your interest. Regardless of which method you have chosen, it is also a good practice to focus on consensus R-loop forming regions based on integrative analysis of existing R-loop mapping datasets (Miller et al. 2021; Lin et al. 2022).

### Theoretical consideration of typical R-loop formation

R-loop formation must enlist nascent RNA transcripts to invade into DNA duplex, especially in genomic regions that contain various non-B DNA structures. There are two models for envisioning this process. The first model considers the topological impact of transcription, which is known to induce positive supercoiling in front of Pol II and negative supercoiling behind. G/C-skewed sequences, which are enriched in gene promoters with high content of CpG islands, coupled with unrelaxed negative supercoiling would favor the formation of non-B DNA structures in the non-template strand to create transient DNA bubbles behind the elongating polymerase. This would enable nascent RNA exiting the RNA channel of the polymerase to re-anneal to the template DNA strand. A free 5’ RNA end would be most efficient to invade into such DNA bubble to gyrate through the duplex DNA, as we recently demonstrated (Chen et al. 2017). This model explains well the frequent formation of R-loops at TSS regions. The free 5’ end generated by the polyadenylation reaction near TTSs may also promote R-loop formation, but the efficiency would be significantly reduced because of both less Pol II pausing and efficient targeting of this unprotected end by the 5’-3’ exonuclease Xrn2 (Skourti-Stathaki et al. 2011). Similarly, R-loop may form with low frequency in enhancer regions as eRNAs are relatively unstable. Importantly, this model also predicts the rare formation of typical R-loop structure of large size (Class 1 in Fig. 6) within gene body because the 5’ RNA end of a nascent RNA may be quickly anchored by assembled complexes, including those for RNA export, thus preventing RNA gyration on the template DNA in a DNA bubble.

The second model would enable the formation of large typical R-loop structures within gene body. In this model, when an elongating RNA polymerase encounters a DNA bubble, it may switch its traveling path from dsDNA to ssDNA, and as a result, the newly synthesized RNA would be able to anneal on the template DNA in a continuous fashion during transcription beyond the bubble. In theory, the model is possible because RNA polymerase is able to transcribe RNA on the ssDNA template in *in vitro* transcription reactions (Hinkle et al. 1972). Inside cells, however, this would require consistent action of Top I to relax supercoiled DNA, which would be consistent with Top I travelling together RNA polymerase during transcription elongation (Baranello et al. 2016). While theoretically possible, it remains unclear how frequently this mode of R-loop formation may occur in cells, a key piece of quantitative information missing from any of the existing mapping strategies. In addition, such gene body signals would be sensitive to RNase H, but insensitive to RNase III, which appears opposite, at least in fission yeast (Hartono et al. 2018). If not efficiently resolved by RNase H, one would also imagine that such large gene body R-loop would block the next round of transcription (Belotserkovskii et al. 2017), which is testable in RNase H inactivated cells. Recently, a similar model is proposed and coined as “RNAP rotation pathway”, and both of us hold that it requires big topological or energy barrier to overcome and thus less likely to happen (Belotserkovskii and Hanawalt 2022). Belotserkovskii and Hanawalt further propose another possibility that disassociation of RNAP might permit the short RNA-DNA hybrid inside the transcription complex to survive for a sufficient period and initiate further RNA invasion into the DNA duplex via strand exchange (Belotserkovskii and Hanawalt 2022). This intriguing model well explains why *ex vivo* assays (e.g., DRIP-seq and DRIPc-seq) but not *in vitro* assays (e.g., R-ChIP) preferentially detect large R-loop formation within gene body, as RNAP disassociation by deproteinization is performed before R-loop detection in *ex vivo* assays (Wang et al. 2021). However, it appears to us that strand exchange *ex vivo* also needs to overcome a kinetic barrier in non-denatured conditions (Crossley et al. 2019). Altogether, although future studies are clearly required to justify our theoretical considerations, we hold that most of the large R-loop peaks detected within gene bodies unlikely correspond to typical R-loop structures.

### Formation and functional implication of two distinct types of RNA:DNA hybrids

An interesting finding in this report is the series of evidence for the second class of RNA:DNA hybrids detected by various current R-loop profiling technologies. We found that this second class of RNA:DNA hybrids is distinguishable from classic R-loops based on different binding patterns of RPA, on both the template and non-template DNA in classic R-loops, but only the template DNA in the second class. In fact, this second class of RNA:DNA hybrids is well known in the DNA replication/repair field. Besides RNA primase-generated RNA on the lagging strand during the DNA replication process, extensive literature has documented DNA damage-induced non-coding RNAs (known as dilncRNAs, which could be further processed into small RNAs by small RNA processing machineries) (Li et al. 2016; Michelini et al. 2017; Cohen et al. 2018; D’Alessandro et al. 2018; Lu et al. 2018). Of note, the current RNase H-based R-loop mapping methods all rely on RNase H1, but RNase H2 is thought to be the major enzyme involved in DNA replication and repair pathways (Williams et al. 2016; Liu et al. 2017; D’Alessandro et al. 2018). However, there is no definitive evidence against the involvement of RNase H1 in these pathways. According to the current model, the 5’ end of broken dsDNA is resected and the exposed 3’ end then serves an ssDNA template for Pol II or Pol III recruited by the MRE11/RAD50/NSB1 complex to transcribe dilncRNAs that can hybridize to the template strand (Liu et al. 2017; D’Alessandro et al. 2018; Liu et al. 2021). This well explains RPA binding on the template strand, as the non-template is removed. Therefore, by definition, this second class of RNA:DNA hybrids is distinct from the three-stranded R-loops.

As DNA damage-induced dilncRNAs can take place anywhere in the genome, it is consistent with our findings that the formation of such RNA:DNA hybrids showed no association with specific histone markers that are normally linked to transcription activities in promoters and enhancers. Intriguingly, our data also showed the lack of nascent RNA signals on these RNA:DNA hybrid-forming regions. One possibility for this apparent discrepancy is that dilncRNA transcription might be somehow disrupted during the GRO-seq and NET-seq procedures. Alternatively, dilncRNAs might be rapidly processed into small RNAs, thus escaping detection by standard nascent RNA mapping technologies. Future studies are required to test these possibilities.

Finally, we wish to point that R-loops have been widely considered to be a threat to the genome, given the tight link between R-loop formation and DNA damage response upon functional inactivation of a critical regulator involved in these processes. However, the cause-consequence relationship is often unclear or even confused. It is conceivable that excessive R-loops may trigger DNA damage, thus representative of a cause to genome instability. On the other hand, dilncRNA transcription followed by DNA repair is clearly a consequence of DNA damage, which may be induced by certain non-B DNA structures in the genome. Because both classes of RNA:DNA hybrids can be detected by immunostaining with S9.6 and DNA damage by γH2AX in cells, neither of these criteria could differentiate between the cause for and consequence of genome instability. Therefore, the accurate assignment of R-loop related RNA:DNA hybrids and DNA damage-induced RNA:DNA hybrids is critical for delineating pathways that lead to genome instability in future studies. Our data provide practical guidelines for distinguishing these two different classes of RNA:DNA hybrids for future mechanistic studies.

## Methods

### Cell lines and cell culture conditions

HEK293T cells were from a common laboratory stock (gift of Dr. Steve Dowdy’s lab of UCSD); K562 cells were purchased from ATCC (ATCC#: CCL-243). HEK293T cells were grown in DMEM supplemented with 10% FBS and 1× penicillin-streptomycin (GIBCO); K562 cells were cultured in RPMI1640 (CORNING) with 10% fetal bovine serum (FBS; Omega Scientific), and 100 U/ml penicillin-streptomycin (GIBCO) at 37 °C in a 5% of CO_2_ incubator.

### Peak calling for different R-loop mapping technologies

DRIP-seq (Manzo et al. 2018), DRIPc-seq (Manzo et al. 2018), RDIP-seq (Nadel et al. 2015), R-ChIP (Chen et al. 2017), R-loop CUT&Tag (Wang et al. 2021) and MapR (Yan et al. 2019) data in HEK293 cells, and ssDRIP-seq (Yang et al. 2019) and RR-ChIP (Tan-Wong et al. 2019) data in Hela cells were downloaded from GEO database. Other technologies were not included for evaluation mainly due to the lack of data in HEK293/Hela cells. Sequencing data from different technical replicates were first combined. All R-loop mapping data were mapped to human genome (hg38) via Bowtie2 (Langmead and Salzberg 2012). Only uniquely-mapped non-redundant sequencing reads (for single-end sequencing data) or read-pairs (for paired-end sequencing data) were kept. Single-end sequencing tags were extended to the estimated average size of sequenced fragments, and the actual fragment size was directly computed for paired-end sequencing data. If existed, control libraries built from input DNA or control treatment were processed as above. To better reflect the signal distribution of individual technologies, broad (DRIP-seq, DRIPc-seq, RDIP-seq, ssDRIP-seq and MapR) or narrow (R-ChIP, R-loop CUT&Tag and RR-ChIP) peak callings were all done with MACS2 for all filtered tags (DRIP-seq and MapR), or filtered tags from Watson or Crick strand separately (DRIPc-seq, RDIP-seq, ssDRIP-seq, R-ChIP and RR-ChIP) (Zhang et al. 2008). Q-value ≤ 0.01, and q-value ≤ 0.001 and fold change ≥ 5 were applied to broad and narrow peaks respectively. When multiple biological replicates were available, we only kept commonly-detected peaks with ≥50% reciprocal overlap between replicates. Human genome was partitioned into 1kb bins, and bins harboring at least one of the top 5,000 R-loop peaks detected by individual technologies and overlapped with GRO-seq peaks in both HEK293 and Hela cells (Chen et al. 2017; Fei et al. 2018) were selected for pairwise comparison (Fig. 1C, E). Bins covered with peaks detected by at least two technologies were considered co-detectable.

### Integrative analysis of public datasets

GRO-seq (Chen et al. 2017; Fei et al. 2018), RPA ChIP-seq (Zhang et al. 2017), GRID-seq (Li et al. 2017), and TOP1-seq (Baranello et al. 2016) were downloaded from GEO database. Processed Repli-seq data were downloaded from ENCODE (Hansen et al. 2010). GRO-seq data were processed and used to check local activity of nascent RNA transcription (Fig. 4A), and to assign the strand information of R-loop peaks detected by non-strand-specific DRIP-seq and MapR wherever needed (Fig. 5C, D). ChIP-seq data for RPA1 and RPA2 were processed as described above. Shared peaks of RPA1 and RPA2 were used for intersection with R-loop peaks detected with different technologies (Fig. 5A). GRID-seq raw data were processed as described previously (Li et al. 2017). Focusing on those Alu elements whose [-5kb, 5kb] regions are fully located within gene body regions, we retrieved GRID-seq RNA tags interacted with its own promoters around Alu elements for comparison with RR-ChIP signals (Fig. 3I). Color space TOP1-seq data were mapped with Bowtie (Langmead et al. 2009). The 5’ ends of uniquely-mapped non-redundant sequencing tags were extracted for representation of TOP1 action sites. Processed PARIS data in HEK293 and Hela cells were directly used for downstream analysis (Lu et al. 2016). Random genomic regions located as the same genes and with the same sizes as those of DRIP(c)-seq or RR-ChIP peaks were generated to compute the expected overlap with PARIS-mapped RNA duplex regions (Fig. 3D). Human A-to-I editing events were collected from REDIportal database (Fig. 3G) (Picardi et al. 2017).

### Annotation of R-loop peaks

Gene models and tRNA loci defined by GENCODE Release 24 was used for genomic annotation (Figs. 1F, 2B). The promoter region was defined as [−1kb, 1kb] from TSS, and terminal region as [−1kb, 1kb] from the poly(A) site, and gene body as the remaining genic region. RepeatMasker-annotated repeat elements were download from UCSC genome browser.

### Detection of Valley(s) within DRIP-seq peaks

For valley detection, base wise DRIP-seq signal intensities measured as reads per million were first computed for any DRIP-seq peaks ≥ 1kb. R package *changepoint* was then applied for step detection (Killick and Eckley 2014). *PELT* method option was set aiming to identify multiple change points if exist. The minimal segment length was set as 200, corresponding to the estimated average DRIP-seq fragment size. Resulting segments showing sufficiently low level of average signals (z-score ≤ -1) were kept, and concatenated if adjacent to each other (Supplemental Fig. S2B). To generate a random set of R-ChIP-mapped R-loops mimicking those overlapped with DRIP-seq peaks, we created regions with the same sizes as those of R-ChIP peaks, and with ≥1bp overlap with corresponding DRIP-seq peaks by Bedtools (Supplemental Fig. S2D) (Quinlan and Hall 2010).

### DRIP-seq with or without RNase H treatment after S9.6 IP in K562 cells

DRIP-seq was performed as described previously (Sanz and Chedin 2019) with some modifications. Briefly, genomic DNA (gDNA) was purified from 8M of K562 cells. The purified gDNA was digested at 37 °C with five restriction enzymes (EcoRI, BsrGI, XbaI, SspI and HindIII from NEB). Ten micrograms of digested gDNA was incubated with 7 μg of S9.6 antibody (Millipore: MABE1095) for IP and the IPed gDNA was recovered by proteinase K (NEB: P8107S) treatment. Before sonication, the purified gDNA was incubated at 37 °C for 1 h with RNase H (Thermo Fisher Scientific: 18021014) to remove hybridized RNA for RNase H-treated DRIP-seq library. DNA fragmented by sonication was repaired using Quick Blunting Kit (NEB: E1201S). For TA ligation, dA was added to the 3’-end of DNA using Klenow fragment exo- (NEB: M0212S) and then the adapter was ligated to the end of DNA. After PCR amplification, DNA was loaded onto acrylamide gel to select library with 200-500 bp in length. DNA library was eluted from the gel fragments for 12 h in elution buffer (10 mM Tris-Cl, pH 8.0, 1 mM EDTA, 0.1% Tween-20, and 300 mM NaCl) and purified using DNA Clean and Concentrator-5 Kit (Zymo Research: D4060).

The libraries were sequenced by HiSeq 4000 system and processed as described above. To test whether valleys were more sensitive to RNase H treatment, a reference set of DRIP-seq peaks and valleys was first detected from a third-part DRIP-seq data in K562 (Sanz et al. 2016). DRIP-seq signal intensities with or without RNase H treatment were then calculated and normalized by subtracting basal levels of signal intensities (Figs. 2E, 3B). *P*-values for whether DRIP-seq signals are increased after RNase H treatment were computed based on Poisson distribution using MACS2 *bdgcmp* sub-command (Fig. 3C) (Zhang et al. 2008).

### Strand-specific RPA ChIP-seq

We took advantage of the ssDNA binding characteristics of RPA protein, and performed strand-specific RPA-ChIP-seq experiment following R-ChIP protocol with modifications (Chen et al. 2019). Briefly, approximately 1×10^7^ HEK293T cells were crosslinked with 1% formaldehyde for 15 min at room temperature. Fixation was stopped by adding Glycine at the final concentration 125 mM followed by incubation on a rotating platform for 15 min at room temperature. After washing plates twice with PBS, cells were scraped off and incubated in cell lysis buffer (10 mM Tris-HCl pH 8.0, 10mM NaCl, 0.5% NP-40 and 1× protease inhibitor cocktail) for 15 min on ice. After centrifugation at 12,000 rpm for 10 min at 4°C, the supernatant was aspirated and discarded.

The cell pellet was then suspended in nuclei lysis buffer (50 mM Tris-HCl pH 8.0, 10 mM EDTA, 1% SDS and 1× protease inhibitor cocktail) and incubated for 10 min on ice. Chromatin DNA was sheared to 250-600 bp in size by sonication. 5% chromatin fragment was saved as input and the remaining was incubated with magnetic beads conjugated with anti-RPA32 antibody (Millipore: EMNA19L-100UG) overnight at 4°C. On the next day, beads were sequentially washed three times with TSEI (20 mM Tris-HCl pH 8.0, 150 mM NaCl, 1% Triton X-100, 0.1% SDS, 2 mM EDTA and 1× protease inhibitor cocktail), three times with TSEII (20 mM Tris-HCl pH 8.0, 500 mM NaCl, 1% Triton X-100, 0.1% SDS, 2 mM EDTA and 1× protease inhibitor cocktail), once with TSEIII (10 mM Tris-HCl pH 8.0, 250 mM LiCl, 1% NP-40, 1% Deoxycholate, 1 mM EDTA and 1× protease inhibitor cocktail) and once with TE buffer (10 mM Tris-HCl pH 8.0 and 1 mM EDTA). The protein-chromatin complex was eluted with elution buffer (10 mM Tris-HCl pH 8.0, 1% SDS and 1 mM EDTA) and reverse-crosslinked on a thermomixer with agitation overnight at 65 °C. After sequential treatments of RNase A and Proteinase K to remove any residual RNA and protein, the precipitated DNA fragment was cleaned twice with phenol and once with phenol:chloroform:isoamyl alcohol, followed by ethanol precipitation. The recovered DNA fragment was subjected to library construction.

To generate the RPA-ChIP-seq library in a strand-specific way, the recovered DNA fragment was subjected to random priming using a primer ended with 9-mer random sequences (5’-/invddt/CAAGCAGAAGACGGCATACGAGNNNNNNNNN-3’). An “A” base was then added to the 3’ end and the adaptor from Illumina was ligated only to one end of the resultant dsDNA as the other end contained a 5’ overhang introduced by the N9 primer. After purification, 16-18 cycles of PCR were performed and PCR products in the size range of 130-350 bp were gel-isolated and purified. Deep sequencing was performed on the Illumina HiSeq 2500 system. The sequenced fragments represent the RPA-bound DNA strand. Strand-specific RPA peaks were called using a similar strategy to R-ChIP (Chen et al. 2019).

## Data access

All raw and processed sequencing data generated in this study have been submitted to the NCBI Gene Expression Omnibus (GEO; https://www.ncbi.nlm.nih.gov/geo/) under accession number GSE147886.

## Competing interest statement

The authors declare no competing financial interests.

## Acknowledgments

We thank members of Fu lab at UCSD and Chen lab at NJU for discussion, and critical reading of the manuscript. This study is supported by NIH grants [HG004659 to X-D.F., DK098808 to X-D.F. and D-E.Z., and DK120952 to J-Y.C.], the Ministry of Science and Technology of China [2021YFF1200017 to J-Y.C. and 2021YFA1100500 to L. C.] and National Natural Science Foundation of China [32170653 to J-Y.C. and 32171289 to L. C.].

## Author contributions

J-Y.C., D-H.L. and X-D.F. conceived the idea. J-Y.C. and Y.Z. analyzed all genomic data. D-H.L. and L.C. performed experiments with help from F.Z., X.Z. and C.S.. H.L. sequenced all libraries. D.W. and D-E.Z. contributed to data interpretation. J-Y.C., D-H.L. and X-D.F. wrote the paper.

## Supplemental Figures and Figure Legends

**Supplemental Figure S1.**
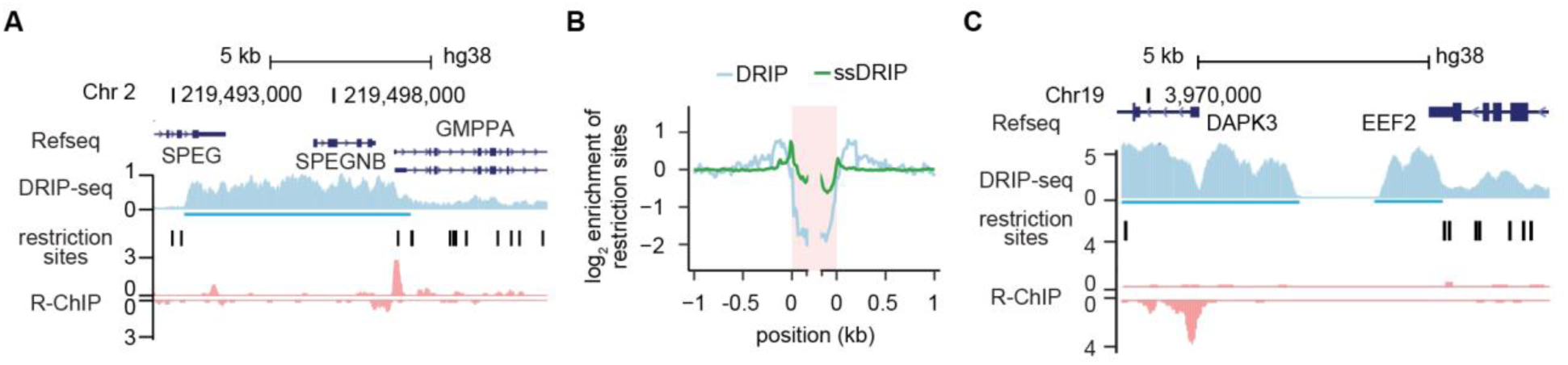
Multi-assignment of broad DRIP-seq peaks to different genomic annotation. (A) A representative broad DRIP-seq peak (light blue underlined) at the *SPEG-*TTS-*SPEGNB*-*GMPPA*-TTS locus in comparison with the restriction sites and strand-specific R-ChIP signals in the region. (B) The ensemble distribution of restriction sites around DRIP-seq and ssDRIP-seq peaks. (C) Two separated peaks, one at *DAPK3*-TSS and the other *EEF2*-TTS region, detected by DRIP-seq within the same restriction fragment where a single R-ChIP peak resides.

**Supplemental Figure S2.**
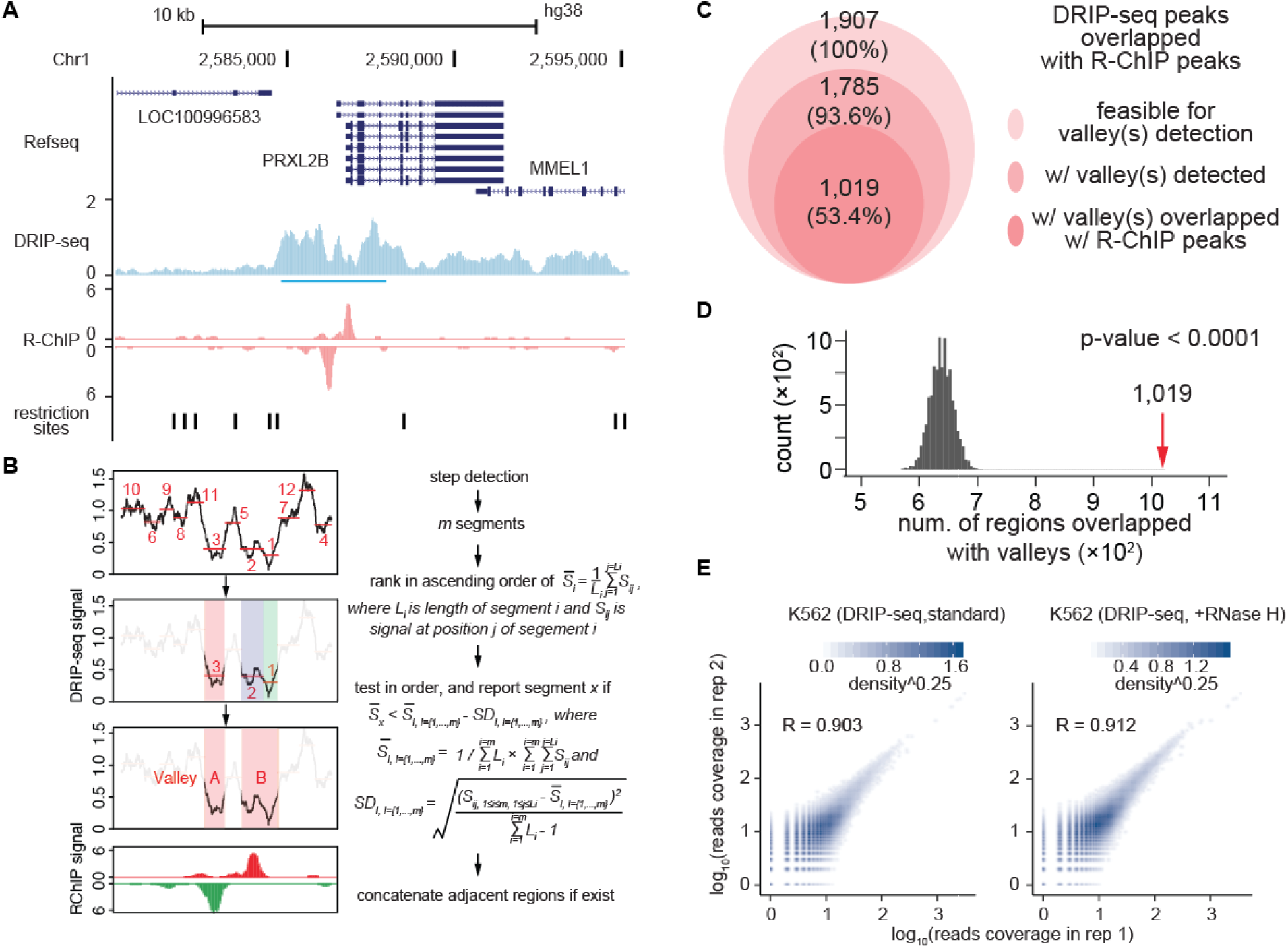
Improvement of DRIP-seq resolution by including an RNase H treatment step. (A) Valleys within a broad DRIP-seq peak in reference to R-ChIP detected peaks. (B) Schematic of the strategy to detect putative valleys within the DRIP-seq peak in (A). (C) Overlap between R-ChIP peaks and detected valleys within DRIP-seq peaks. The analysis began with a set of commonly detected peaks (light pink circle), from which valleys were identified according to the strategy in B (medium pink circle) followed by matching valleys in DRIP-seq peaks with R-ChIP peaks (dark pink circle). (D) Distribution of the expected frequency of overlaps between simulated R-ChIP peaks and detected valleys after 10,000-time simulations. Red arrow marks the observed overlap, which happens less than once after 10,000-time simulations (*p*-value < 0.0001). (E) Reproducibility of duplicated DRIP-seq libraries with or without RNase H treatment, as indicated by Pearson’s correlation coefficients.

**Supplemental Figure S3.**
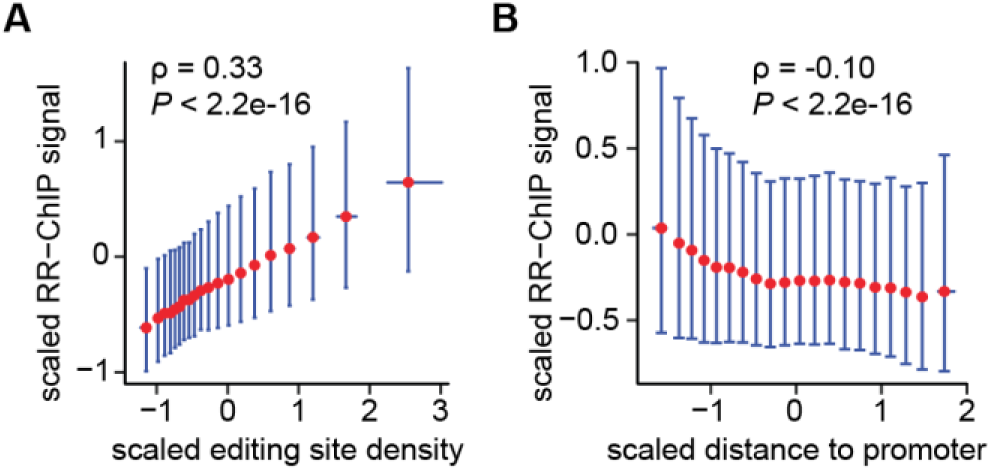
Evaluation of R-loop peaks within gene body regions. (A) Correlation of RR-ChIP-captured signals and the propensity to form structured RNA, as measured by RNA editing site density at Alu repeats. Data are shown as median (red dot) ± [25%,75%] intervals. RR-ChIP signals and editing site density (site per bp) are scaled for each gene. Spearman correlation (ρ) and *p*-value are shown. (B) Correlation of RR-ChIP-captured signals on Alu repeats and their distances to promoter regions. Data are shown as median (red dot) ± [25%,75%] intervals. RR-ChIP signals and distances to promoter regions are scaled for each gene. Spearman correlation (ρ) and *p*-value are shown.

**Supplemental Figure S4.**
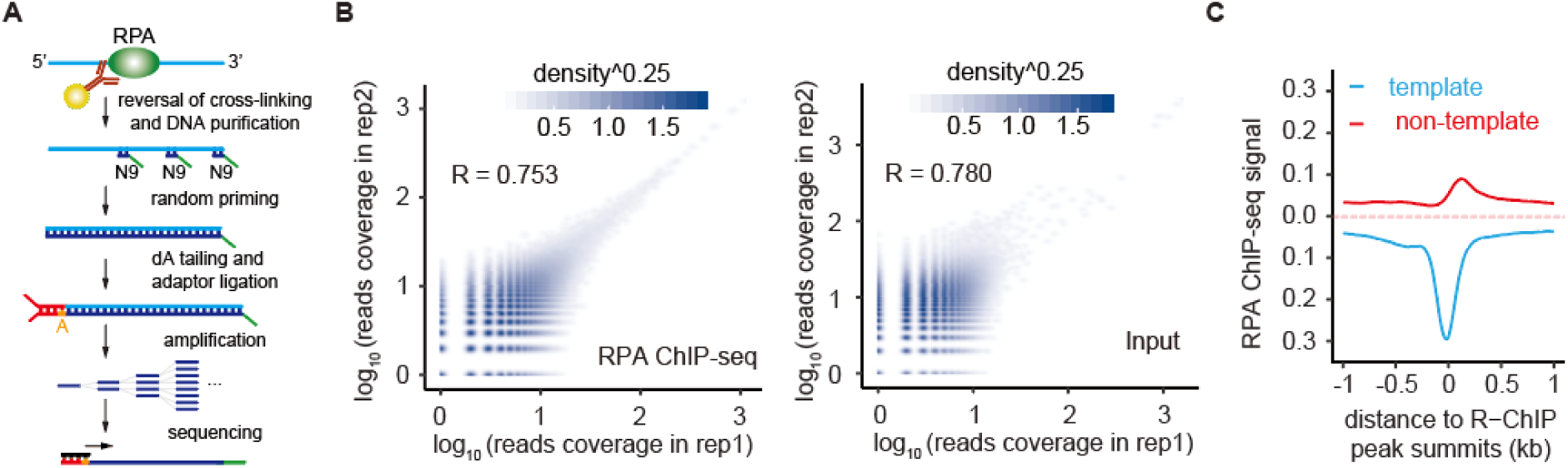
Evaluation of RPA ChIP-seq data generated in HEK293 cells. (A) Procedure of strand-specific RPA ChIP-seq. (B) Pearson’s correlation between duplicated RPA ChIP-seq (left) and input (right) libraries. (C) Alignment of strand-specific RPA ChIP-seq signals on R-loop peaks with no antisense R-loop peaks around.

## Notes

### Competing Interest Statement

The authors have declared no competing interest.

## References

Arab K, Karaulanov E, Musheev M, Trnka P, Schafer A, Grummt I, Niehrs C. 2019. GADD45A binds R-loops and recruits TET1 to CpG island promoters. Nat Genet 51: 217–223.

Bader AS, Hawley BR, Wilczynska A, Bushell M. 2020. The roles of RNA in DNA double-strand break repair. Br J Cancer 122: 613–623.

Bai X, Li F, Zhang Z. 2021. A hypothetical model of trans-acting R-loops-mediated promoter-enhancer interactions by Alu elements. J Genet Genomics 48: 1007–1019.

Baranello L, Wojtowicz D, Cui K, Devaiah BN, Chung HJ, Chan-Salis KY, Guha R, Wilson K, Zhang X, Zhang H et al. 2016. RNA Polymerase II Regulates Topoisomerase 1 Activity to Favor Efficient Transcription. Cell 165: 357–371.

Baranovskiy AG, Babayeva ND, Zhang Y, Gu J, Suwa Y, Pavlov YI, Tahirov TH. 2016. Mechanism of Concerted RNA-DNA Primer Synthesis by the Human Primosome. J Biol Chem 291: 10006–10020.

Belotserkovskii BP, Hanawalt PC. 2022. Mechanism for R-loop formation remote from the transcription start site: Topological issues and possible facilitation by dissociation of RNA polymerase. DNA Repair (Amst) 110: 103275.

Belotserkovskii BP, Soo Shin JH, Hanawalt PC. 2017. Strong transcription blockage mediated by R-loop formation within a G-rich homopurine-homopyrimidine sequence localized in the vicinity of the promoter. Nucleic Acids Res 45: 6589–6599.

Castillo-Guzman D, Chedin F. 2021. Defining R-loop classes and their contributions to genome instability. DNA Repair (Amst) 106: 103182.

Chen JY, Zhang X, Fu XD, Chen L. 2019. R-ChIP for genome-wide mapping of R-loops by using catalytically inactive RNASEH1. Nat Protoc 14: 1661–1685.

Chen L, Chen JY, Zhang X, Gu Y, Xiao R, Shao C, Tang P, Qian H, Luo D, Li H et al. 2017. R-ChIP Using Inactive RNase H Reveals Dynamic Coupling of R-loops with Transcriptional Pausing at Gene Promoters. Mol Cell 68: 745–757 e745.

Chen PB, Chen HV, Acharya D, Rando OJ, Fazzio TG. 2015. R loops regulate promoter-proximal chromatin architecture and cellular differentiation. Nat Struct Mol Biol 22: 999–1007.

Cohen S, Puget N, Lin YL, Clouaire T, Aguirrebengoa M, Rocher V, Pasero P, Canitrot Y, Legube G. 2018. Senataxin resolves RNA:DNA hybrids forming at DNA double-strand breaks to prevent translocations. Nat Commun 9: 533.

Crossley MP, Bocek M, Cimprich KA. 2019. R-Loops as Cellular Regulators and Genomic Threats. Mol Cell 73: 398–411.

Crossley MP, Bocek MJ, Hamperl S, Swigut T, Cimprich KA. 2020. qDRIP: a method to quantitatively assess RNA-DNA hybrid formation genome-wide. Nucleic Acids Res 48: e84.

D’Alessandro G, Whelan DR, Howard SM, Vitelli V, Renaudin X, Adamowicz M, Iannelli F, Jones-Weinert CW, Lee M, Matti V et al. 2018. BRCA2 controls DNA:RNA hybrid level at DSBs by mediating RNase H2 recruitment. Nat Commun 9: 5376.

Dumelie JG, Jaffrey SR. 2017. Defining the location of promoter-associated R-loops at near-nucleotide resolution using bisDRIP-seq. Elife 6: e28306.

Duquette ML, Handa P, Vincent JA, Taylor AF, Maizels N. 2004. Intracellular transcription of G-rich DNAs induces formation of G-loops, novel structures containing G4 DNA. Genes Dev 18: 1618–1629.

El Hage A, Webb S, Kerr A, Tollervey D. 2014. Genome-wide distribution of RNA-DNA hybrids identifies RNase H targets in tRNA genes, retrotransposons and mitochondria. PLoS Genet 10: e1004716.

Fei J, Ishii H, Hoeksema MA, Meitinger F, Kassavetis GA, Glass CK, Ren B, Kadonaga JT. 2018. NDF, a nucleosome-destabilizing factor that facilitates transcription through nucleosomes. Genes Dev 32: 682–694.

Gan W, Guan Z, Liu J, Gui T, Shen K, Manley JL, Li X. 2011. R-loop-mediated genomic instability is caused by impairment of replication fork progression. Genes Dev 25: 2041–2056.

Garcia-Muse T, Aguilera A. 2019. R Loops: From Physiological to Pathological Roles. Cell 179: 604–618.

Ginno PA, Lott PL, Christensen HC, Korf I, Chedin F. 2012. R-loop formation is a distinctive characteristic of unmethylated human CpG island promoters. Mol Cell 45: 814–825.

Halasz L, Karanyi Z, Boros-Olah B, Kuik-Rozsa T, Sipos E, Nagy E, Mosolygo LA, Mazlo A, Rajnavolgyi E, Halmos G et al. 2017. RNA-DNA hybrid (R-loop) immunoprecipitation mapping: an analytical workflow to evaluate inherent biases. Genome Res 27: 1063–1073.

Hansen RS, Thomas S, Sandstrom R, Canfield TK, Thurman RE, Weaver M, Dorschner MO, Gartler SM, Stamatoyannopoulos JA. 2010. Sequencing newly replicated DNA reveals widespread plasticity in human replication timing. Proc Natl Acad Sci U S A 107: 139–144.

Hartono SR, Malapert A, Legros P, Bernard P, Chedin F, Vanoosthuyse V. 2018. The Affinity of the S9.6 Antibody for Double-Stranded RNAs Impacts the Accurate Mapping of R-Loops in Fission Yeast. J Mol Biol 430: 272–284.

Hawley BR, Lu WT, Wilczynska A, Bushell M. 2017. The emerging role of RNAs in DNA damage repair. Cell Death Differ 24: 580–587.

Hinkle DC, Ring J, Chamberlin MJ. 1972. Studies of the binding of Escherichia coli RNA polymerase to DNA. 3. Tight binding of RNA polymerase holoenzyme to single-strand breaks in T7 DNA. J Mol Biol 70: 197–207.

Jauregui-Lozano J, Escobedo S, Easton A, Lanman NA, Weake VM, Hall H. 2022. Proper control of R-loop homeostasis is required for maintenance of gene expression and neuronal function during aging. Aging Cell 21: e13554.

Jimeno S, Prados-Carvajal R, Fernandez-Avila MJ, Silva S, Silvestris DA, Endara-Coll M, Rodriguez-Real G, Domingo-Prim J, Mejias-Navarro F, Romero-Franco A et al. 2021. ADAR-mediated RNA editing of DNA:RNA hybrids is required for DNA double strand break repair. Nat Commun 12: 5512.

Kabeche L, Nguyen HD, Buisson R, Zou L. 2018. A mitosis-specific and R loop-driven ATR pathway promotes faithful chromosome segregation. Science 359: 108–114.

Killick R, Eckley IA. 2014. changepoint: An R Package for Changepoint Analysis. Journal of Statistical Software 58: 1–19.

Langmead B, Salzberg SL. 2012. Fast gapped-read alignment with Bowtie 2. Nat Methods 9: 357–359.

Langmead B, Trapnell C, Pop M, Salzberg SL. 2009. Ultrafast and memory-efficient alignment of short DNA sequences to the human genome. Genome Biol 10: R25.

Li L, Germain DR, Poon HY, Hildebrandt MR, Monckton EA, McDonald D, Hendzel MJ, Godbout R. 2016. DEAD Box 1 Facilitates Removal of RNA and Homologous Recombination at DNA Double-Strand Breaks. Mol Cell Biol 36: 2794–2810.

Li X, Zhou B, Chen L, Gou LT, Li H, Fu XD. 2017. GRID-seq reveals the global RNA-chromatin interactome. Nat Biotechnol 35: 940–950.

Lin R, Zhong X, Zhou Y, Geng H, Hu Q, Huang Z, Hu J, Fu XD, Chen L, Chen JY. 2022. R-loopBase: a knowledgebase for genome-wide R-loop formation and regulation. Nucleic Acids Res 50: D303–D315.

Liu B, Hu J, Wang J, Kong D. 2017. Direct Visualization of RNA-DNA Primer Removal from Okazaki Fragments Provides Support for Flap Cleavage and Exonucleolytic Pathways in Eukaryotic Cells. J Biol Chem 292: 4777–4788.

Liu S, Hua Y, Wang J, Li L, Yuan J, Zhang B, Wang Z, Ji J, Kong D. 2021. RNA polymerase III is required for the repair of DNA double-strand breaks by homologous recombination. Cell 184: 1314–1329 e1310.

Lu WT, Hawley BR, Skalka GL, Baldock RA, Smith EM, Bader AS, Malewicz M, Watts FZ, Wilczynska A, Bushell M. 2018. Drosha drives the formation of DNA:RNA hybrids around DNA break sites to facilitate DNA repair. Nat Commun 9: 532.

Lu Z, Zhang QC, Lee B, Flynn RA, Smith MA, Robinson JT, Davidovich C, Gooding AR, Goodrich KJ, Mattick JS et al. 2016. RNA Duplex Map in Living Cells Reveals Higher-Order Transcriptome Structure. Cell 165: 1267–1279.

Manzo SG, Hartono SR, Sanz LA, Marinello J, De Biasi S, Cossarizza A, Capranico G, Chedin F. 2018. DNA Topoisomerase I differentially modulates R-loops across the human genome. Genome Biol 19: 100.

Marechal A, Zou L. 2015. RPA-coated single-stranded DNA as a platform for post-translational modifications in the DNA damage response. Cell Res 25: 9–23.

Mayer A, di Iulio J, Maleri S, Eser U, Vierstra J, Reynolds A, Sandstrom R, Stamatoyannopoulos JA, Churchman LS. 2015. Native elongating transcript sequencing reveals human transcriptional activity at nucleotide resolution. Cell 161: 541–554.

Michelini F, Pitchiaya S, Vitelli V, Sharma S, Gioia U, Pessina F, Cabrini M, Wang Y, Capozzo I, Iannelli F et al. 2017. Damage-induced lncRNAs control the DNA damage response through interaction with DDRNAs at individual double-strand breaks. Nat Cell Biol 19: 1400–1411.

Miller HE, Montemayor D, Abdul J, Vines A, Levy S, Hartono S, Sharma K, Frost B, Chedin F, Bishop AJR. 2021. Quality-controlled R-loop meta-analysis reveals the characteristics of R-Loop consensus regions. BioRxiv doi:10.1101/2021.11.01.466823.

Nadel J, Athanasiadou R, Lemetre C, Wijetunga NA, P OB, Sato H, Zhang Z, Jeddeloh J, Montagna C, Golden A et al. 2015. RNA:DNA hybrids in the human genome have distinctive nucleotide characteristics, chromatin composition, and transcriptional relationships. Epigenetics Chromatin 8: 46.

Nguyen HD, Yadav T, Giri S, Saez B, Graubert TA, Zou L. 2017. Functions of Replication Protein A as a Sensor of R Loops and a Regulator of RNaseH1. Mol Cell 65: 832–847 e834.

Nishikura K. 2010. Functions and regulation of RNA editing by ADAR deaminases. Annu Rev Biochem 79: 321–349.

Nowotny M, Cerritelli SM, Ghirlando R, Gaidamakov SA, Crouch RJ, Yang W. 2008. Specific recognition of RNA/DNA hybrid and enhancement of human RNase H1 activity by HBD. EMBO J 27: 1172–1181.

Ohle C, Tesorero R, Schermann G, Dobrev N, Sinning I, Fischer T. 2016. Transient RNA-DNA Hybrids Are Required for Efficient Double-Strand Break Repair. Cell 167: 1001–1013 e1007.

Ouyang J, Yadav T, Zhang JM, Yang H, Rheinbay E, Guo H, Haber DA, Lan L, Zou L. 2021. RNA transcripts stimulate homologous recombination by forming DR-loops. Nature 594: 283–288.

Picardi E, D’Erchia AM, Lo Giudice C, Pesole G. 2017. REDIportal: a comprehensive database of A-to-I RNA editing events in humans. Nucleic Acids Res 45: D750–D757.

Quinlan AR, Hall IM. 2010. BEDTools: a flexible suite of utilities for comparing genomic features. Bioinformatics 26: 841–842.

Richard P, Manley JL. 2017. R Loops and Links to Human Disease. J Mol Biol 429: 3168–3180.

Roberts RW, Crothers DM. 1992. Stability and properties of double and triple helices: dramatic effects of RNA or DNA backbone composition. Science 258: 1463–1466.

Rouillard AD, Gundersen GW, Fernandez NF, Wang Z, Monteiro CD, McDermott MG, Ma’ayan A. 2016. The harmonizome: a collection of processed datasets gathered to serve and mine knowledge about genes and proteins. Database (Oxford) 2016.

Sanz LA, Chedin F. 2019. High-resolution, strand-specific R-loop mapping via S9.6-based DNA-RNA immunoprecipitation and high-throughput sequencing. Nat Protoc 14: 1734–1755.

Sanz LA, Hartono SR, Lim YW, Steyaert S, Rajpurkar A, Ginno PA, Xu X, Chedin F. 2016. Prevalent, Dynamic, and Conserved R-Loop Structures Associate with Specific Epigenomic Signatures in Mammals. Mol Cell 63: 167–178.

Shin SI, Ham S, Park J, Seo SH, Lim CH, Jeon H, Huh J, Roh TY. 2016. Z-DNA-forming sites identified by ChIP-Seq are associated with actively transcribed regions in the human genome. DNA Res 23: 477–486.

Skourti-Stathaki K, Proudfoot NJ, Gromak N. 2011. Human senataxin resolves RNA/DNA hybrids formed at transcriptional pause sites to promote Xrn2-dependent termination. Mol Cell 42: 794–805.

Sollier J, Stork CT, Garcia-Rubio ML, Paulsen RD, Aguilera A, Cimprich KA. 2014. Transcription-coupled nucleotide excision repair factors promote R-loop-induced genome instability. Mol Cell 56: 777–785.

Stamatoyannopoulos JA, Adzhubei I, Thurman RE, Kryukov GV, Mirkin SM, Sunyaev SR. 2009. Human mutation rate associated with DNA replication timing. Nat Genet 41: 393–395.

Tan-Wong SM, Dhir S, Proudfoot NJ. 2019. R-Loops Promote Antisense Transcription across the Mammalian Genome. Mol Cell 76: 600–616 e606.

Vanoosthuyse V. 2018. Strengths and Weaknesses of the Current Strategies to Map and Characterize R-Loops. Noncoding RNA 4: 9.

Wahba L, Costantino L, Tan FJ, Zimmer A, Koshland D. 2016. S1-DRIP-seq identifies high expression and polyA tracts as major contributors to R-loop formation. Genes Dev 30: 1327–1338.

Wang K, Wang H, Li C, Yin Z, Xiao R, Li Q, Xiang Y, Wang W, Huang J, Chen L et al. 2021. Genomic profiling of native R loops with a DNA-RNA hybrid recognition sensor. Sci Adv 7.

Williams JS, Lujan SA, Kunkel TA. 2016. Processing ribonucleotides incorporated during eukaryotic DNA replication. Nat Rev Mol Cell Biol 17: 350–363.

Wulfridge P, Sarma K. 2021. A nuclease- and bisulfite-based strategy captures strand-specific R-loops genome-wide. Elife 10.

Xu W, Xu H, Li K, Fan Y, Liu Y, Yang X, Sun Q. 2017. The R-loop is a common chromatin feature of the Arabidopsis genome. Nat Plants 3: 704–714.

Yan Q, Shields EJ, Bonasio R, Sarma K. 2019. Mapping Native R-Loops Genome-wide Using a Targeted Nuclease Approach. Cell Rep 29: 1369–1380 e1365.

Yang X, Liu QL, Xu W, Zhang YC, Yang Y, Ju LF, Chen J, Chen YS, Li K, Ren J et al. 2019. m(6)A promotes R-loop formation to facilitate transcription termination. Cell Res 29: 1035–1038.

Zenkin N, Naryshkina T, Kuznedelov K, Severinov K. 2006. The mechanism of DNA replication primer synthesis by RNA polymerase. Nature 439: 617–620.

Zhang H, Gan H, Wang Z, Lee JH, Zhou H, Ordog T, Wold MS, Ljungman M, Zhang Z. 2017. RPA Interacts with HIRA and Regulates H3.3 Deposition at Gene Regulatory Elements in Mammalian Cells. Mol Cell 65: 272–284.

Zhang Y, Liu T, Meyer CA, Eeckhoute J, Johnson DS, Bernstein BE, Nusbaum C, Myers RM, Brown M, Li W et al. 2008. Model-based analysis of ChIP-Seq (MACS). Genome Biol 9: R137.

Zheng Y, Lorenzo C, Beal PA. 2017. DNA editing in DNA/RNA hybrids by adenosine deaminases that act on RNA. Nucleic Acids Res 45: 3369–3377.

